# An optimized framework for simultaneous EEG-fMRI at 7T enabling safe, high-quality human brain imaging with millisecond temporal resolution and sub-millimeter spatial resolution

**DOI:** 10.1101/2024.08.23.609301

**Authors:** Cristina Sainz Martinez, Jonathan Wirsich, Cilia Jäger, Tracy Warbrick, Serge Vulliémoz, Mathieu Lemay, Jessica Bastiaansen, Roland Wiest, João Jorge

## Abstract

The combination of electroencephalography (EEG) and functional magnetic resonance imaging (fMRI) at ultra-high field (7 Tesla) offers appealing new possibilities to probe human brain function non-invasively with high coverage, millisecond temporal precision and sub-millimeter spatial precision, unraveling cortical layers and small subcortical structures. Unfortunately, this technique has remained largely inaccessible at 7T, due to prohibitive cross-modal interference effects and physical constraints. Here, we developed a first-of-its-kind EEG-fMRI acquisition framework on a clinical 7T system combining key improvements from previous research: compact EEG transmission to reduce artifact incidence, reference sensors for artifact correction, and adapted leads for compatibility with a dense radiofrequency receive-array allowing state-of-the-art fMRI sensitivity and acceleration. Two implementations were tested: one using an EEG cap adapted in-house, and another using a recently-designed prototype from an industrial manufacturer, intended to be further developed into a commercial device accessible to the broader community. A comprehensive evaluation in humans showed that simultaneous acquisitions, including sub-millimeter fMRI resolution, could be conducted without detectable safety issues or major practical constraints. The EEG exerted relatively mild perturbations on fMRI quality (6–11% loss in temporal SNR), without measurably affecting the detection of resting-state networks and visual responses. The artifacts induced on EEG could be corrected to a degree where the spatial, spectral and temporal characteristics were comparable to outside recordings, and hallmark features such as resting-state alpha and eyes-closing alpha modulation could be clearly detected. Altogether, these findings indicate excellent prospects for neuroimaging applications, that can leverage the unique possibilities achievable at 7T.

## 1. Introduction

The neuronal, vascular and metabolic processes that orchestrate brain function are remarkably complex and diverse (Iadecola, 2017; Schaeffer and Iadecola, 2021), and call for more powerful neuroimaging approaches that can combine information from multiple modalities, to achieve more complete pictures of the mechanisms at play (Uludağ and Roebroeck, 2014). In this context, the combination of non-invasive electroencephalography (EEG) and functional magnetic resonance imaging (fMRI) has proved particularly attractive, as the former technique measures the electric field fluctuations generated by neuronal post-synaptic activity, with high temporal specificity, while the latter captures the accompanying hemodynamic fluctuations, with higher spatial specificity (Jorge et al., 2014). When acquired simultaneously, EEG and fMRI can probe highly-relevant spontaneously occurring processes, such as the generation and propagation of epileptic activity (van Graan et al., 2015), interactions within and across brain networks (Sadaghiani and Wirsich, 2020), and more recently, interactions between neuronal activity, hemodynamics and waste clearance during sleep (Fultz et al., 2019), which may play a crucial role in neurodegenerative processes (Hanke et al., 2022).

A key line of development in fMRI has been the pursuit of higher magnetic field strengths, which offer not only higher signal-to-noise ratio (SNR) (Le Ster et al., 2022), but also strong sensitivity boosts specifically for fMRI (Van Der Zwaag et al., 2009). In particular, the sensitivity achieved at 7T has allowed reliable human acquisitions at sub-millimeter resolution, enabling the study of function at the level of cortical columns (Yacoub et al., 2008) and layers (Polimeni et al., 2010), and more specific inferences on the directionality of interactions (Yang et al., 2021). Likewise, the combination of 7T fMRI with EEG offers unprecedented opportunities for basic and clinical neuroscience: for example, in investigating layer-specific activity associated to intra-cortical and thalamocortical communication (Scheeringa and Fries, 2019), or in mapping epileptic sources more precisely (Grouiller et al., 2016), of crucial importance to presurgical planning in pharmaco-resistant patients (Thornton et al., 2011).

Unfortunately, when combined, EEG and fMRI exert important electromagnetic interferences on each other, which introduce specific safety risks for the patients (Dempsey et al., 2001) and EEG equipment (Jorge et al., 2015b), and can cause substantial artifacts on the data (Mullinger and Bowtell, 2011). Crucially, these effects become increasingly more challenging at higher MR field strengths (Mullinger et al., 2008b; Neuner et al., 2014), and the resulting artifacts have strongly limited applications at 7T (Bonmassar et al., 2022). These challenges have motivated extensive research efforts over the last two decades, including advances in acquisition and in data processing (Warbrick, 2022). Seminal studies focused on subject safety proposed hardware additions such as safety resistors on the EEG leads (Lemieux et al., 1997), and identified important aspects to control in MRI sequences (Nöth et al., 2012). The foremost challenge was then the quality of EEG data, affected by artifacts that can be several orders of magnitude larger than the signals of interest (Mullinger and Bowtell, 2011). This includes contributions from the time-varying MRI gradient fields used for image encoding (typically termed “gradient artifacts” (GA)) (Yan et al., 2009), head motion and blood pulsation linked to cardiac activity (termed ballistocardiogram or “pulse artifacts” (PA)) (Yan et al., 2010), spontaneous motion of the head in the static field B_0_ (termed “motion artifacts” (MA)) (Jorge et al., 2015a), and perturbations from other modules of the MRI scanner (Nierhaus et al., 2013; Rothlübbers et al., 2015) (hereafter termed “environment artifacts” (EA)). Numerous studies have proposed post-processing methods to address these different contributions, leveraging model-based (Allen et al., 2000, 1998) and data-based approaches such as independent component analysis (ICA) (Abreu et al., 2018). On the acquisition side, landmark advances included the pursuit of more compact EEG recording setups, with shorter and better-shielded transmission leads to reduce the incidence of artifacts at their origin (Assecondi et al., 2016; Jorge et al., 2015b), and the introduction of dedicated sensors to monitor EEG artifacts, to then correct the EEG channels (Chowdhury et al., 2014; LeVan et al., 2013; Masterton et al., 2007).

With EEG artifacts becoming better accounted for, the penalties on the MRI side have become an increasingly pertinent focus of research. Here, the most important obstacles have been found to be (i) the disruption of MR radiofrequency (RF) pulses, also known as B_1_ field, by the presence of EEG components on the scalp (Angelone et al., 2004), and (ii) physical constraints in the placement and routing of EEG equipment in the confined space of high-density head RF arrays. Regarding the first problem, insights from dedicated studies suggest that B_1_ disruption may be caused predominantly by RF interactions with the conductive EEG leads (Hawsawi et al., 2017; Lê et al., 2022), which become particularly problematic at 7T and above (Dempsey et al., 2001). To mitigate these interactions, some studies have proposed the use of more resistive materials for the leads (Poulsen et al., 2017; Vasios et al., 2006), or the addition of resistors to segment their lengths (Lê et al., 2022). Although showing promise in terms of B_1_ preservation, the ideas proposed so far have unfortunately presented practical obstacles for scalable implementation and usage, which still lack definitive solutions.

The obstacle of physical constraints inside the RF coil also tends to become more problematic at higher fields: to mitigate the associated decrease in RF tissue penetration (Röschmann, 1987), and to accelerate image acquisitions via parallel imaging (Setsompop et al., 2016), the best RF arrays currently available are designed in dense “meshes” of small receive elements, closely covering the head (Hernandez and Kim, 2020). This affords high SNR and acceleration flexibility at 7T, and is currently the preferred design for fMRI at sub-millimeter resolution (Feinberg et al., 2023). Unfortunately, it also imposes strong constraints for fitting EEG caps and routing their leads to exit the coil space and connect to amplifiers. Initial EEG-fMRI studies at 7T were more focused on EEG quality, and thus conceded to using more spacious, though less powerful, RF coils (Jorge et al., 2015b; Mullinger et al., 2008b) – therein undercutting the fMRI gains offered by 7T. More recently, an insightful user modification of a commercial EEG cap model was shown to allow its use together with one of the most widely adopted commercial RF arrays at 7T (Meyer et al., 2020). With that setup, the authors were able to acquire simultaneous EEG-fMRI and extract meaningful features from both modalities. Curiously, the MRI data also appeared to experience milder degradation effects than observed with previous setups at 7T (Jorge et al., 2015b; Mullinger et al., 2008b). Unfortunately, the absence of direct comparisons of MRI data acquired with and without EEG, in that work, prevented more quantitative insights into the extent of those EEG-induced losses in SNR and functional sensitivity (Meyer et al., 2020).

As human 7T systems become increasingly available across the world, enabling unprecedented insights into fine-scale brain function and dysfunction, the interest in reliable frameworks to combine EEG with 7T fMRI is also rapidly increasing. To meet this need, it is now crucial to harness together and improve upon the most promising technological ideas proposed in the literature, to comprehensively assess their safety and effectiveness, and to implement robust setups that can be widely distributed and used by scientists and clinicians of broad background. In the present work, we have sought to address these critical needs. We developed a new acquisition framework for 7T EEG-fMRI combining, for the first time, three landmark improvements: (i) compact EEG setup with short transmission leads between the cap and amplifiers, to reduce the incidence of induction artifacts (Jorge et al., 2015b), (ii) reference sensors to capture artifacts and help denoising the EEG signals (Jorge et al., 2015a; Masterton et al., 2007; Van Der Meer et al., 2016), and (iii) adapted EEG transmission chain with slim cables to make the setup compatible with a state-of-the-art dense receive RF coil. Two implementations of this framework were tested: one based on an EEG cap modified in-house, and another using a recently-designed prototype from an industrial manufacturer (BrainCap MR7Flex) – both tailored for this application. The framework was extensively tested on a group of healthy volunteers, including a sub-millimeter fMRI protocol, to perform a comprehensive evaluation of safety, data quality and functional sensitivity (resting-state activity and eyes-open/closed responses) on both modalities.

## 2. Methods

This study was approved by the local ethics committee (KEK Bern) and by the Swiss regulatory authority for medical devices (Swissmedic), and included a total of 17 healthy adults, sub-divided in two groups, who provided written informed consent before participating. The first group (N=8, 4 male/4 female, 28±4 years old) underwent extensive MRI tests with and without concurrent EEG, to allow a direct evaluation of the impact of the EEG system on MRI data quality and functional sensitivity. The second group (N=9, 5 male/4 female, 23±1 years old) underwent only EEG recordings, outside the MRI environment, to serve as a reference for EEG quality and spatiotemporal characteristics, for comparison to the EEG data acquired concurrently with fMRI in the first group.

### 2.1. Data acquisition

Prior to human scans, the EEG-MRI setup (with both caps) was tested on an agar phantom with concurrent temperature monitoring, to assess safety. The first human scans also included temperature monitoring. The first participant group underwent a 7T MRI session including field mapping, structural and functional acquisitions (detailed below). For each participant, the acquisitions were first performed without, and then repeated with, concurrent EEG (in-house cap prototype). From this group, a subset of N=5 (selected simply based on availability) returned at a later date to repeat the full protocol without and with the BrainCap MR7Flex. In each visit, the acquisitions without EEG were performed first, to avoid gel residues on the head. All human fMRI runs included pulse and breathing recordings (using the default physiological monitoring unit of the scanner), but these were not analyzed in this work.

#### 2.1.1. MRI system and sequences

MRI was performed with a CE-marked 7T Magnetom Terra system (Siemens Healthineers AG, Erlangen, Germany), equipped with XR gradients (80 mT/m at 200 T/m/s) and a single-channel transmit, 32-channel receive head RF coil (Nova Medical, Wilmington MA, USA). The acquisition protocol (detailed in Supp. Table I) included a low-SAR structural scan (3D-GRE), a T_1_-weighted structural (3D-MP2RAGE), a B_0_ map (double-echo 2D-GRE), a B_1_^+^ map (3D-SA2RAGE), fMRI runs at 1.6 mm and at 0.8 mm resolution (2D SMS GE-EPI), and reference EPI scans (2D SE-EPI) matched to the fMRI to guide distortion correction. All protocols covered the whole brain, except for the B_0_ and 0.8 mm fMRI sequences, which excluded a part of the cerebellum and lower regions.

The B_1_^+^ root-mean-square (B_1_^+^_RMS_) (Egan et al., 2021) and specific absorption rate (SAR) were saved from the RF logs of every acquisition. The patient ventilation and lighting were switched off, whereas the Helium coldheads functioned as normal. In this study, the MP2RAGE and SE-EPI sequences were only tested without EEG in the human group, but were also tested with EEG on the phantom, together with temperature monitoring: SE-EPI was tested as a means to push RF energy deposition further and evaluate the extent and location of potentially stronger heating effects, to better understand what may occur when scanning at higher SAR conditions; the MP2RAGE was added to investigate the possibility of performing the T_1_-weighted structural scan as part of the main EEG-MRI session, which could prove convenient to future studies to avoid needing a separate, MRI-only run.

For each participant, at the start of the no-EEG acquisitions, the RF transmit voltage proposed by the scanner calibrations (“B1map 2D / AdjTra2D” procedure) was noted, and fixed for all acquisitions without EEG (hereafter termed V_ref_). In the following acquisitions with EEG, a new transmit value was estimated by the same procedure (hereafter termed V_adj_). To better evaluate the impact of the EEG on image quality, without the confound of transmit voltage compensation, the B_1_ ^+^ mapping and structural GRE acquisitions with EEG were performed once with V_ref_, and another time with V_adj_. The other protocols (B_0_, fMRI) were only performed using V_adj_.

#### 2.1.2. EEG system

The EEG recording setup was tested in two variants (Fig. 1): one using an adaptation of a commercial cap developed in-house, and the other using a prototype developed by Brain Products GmbH (Munich, Germany), designated BrainCap MR7Flex. Both derived from a more standard BrainCap MR model (EasyCap, Herrsching, Germany, Fig. 1-1), with 64 sintered Ag/AgCl ring-type electrodes arranged according to the 10-20 system, with one electrode extended to the back for ECG recording. Each electrode connected to a 5 kΩ current-limiting resistor followed by a standard tinsel lead, which then interfaced with more specific adaptations:

- **In-house prototype:** Similar to the approach of (Meyer et al., 2020), the EEG cap leads were routed towards the parietal area, and soldered to flat ribbon cables that can fit through the narrow gap between the two sliding halves of the Nova RF coil (Fig. 1-4,2). Differently from (Meyer et al., 2020), we reduced exposed areas by employing two ribbon cables (34 tracks each) stacked one on top of the other, which can fit through the coil gap. After exiting the coil, the ribbon cable tracks were re-soldered to tinsel leads, bundled and terminated in custom compact connectors to the EEG amplifiers (Fig. 1-3). Altogether, the distance between the top of the cap (Fig. 1-5) and the amplifier connection was approximately 37 cm. During the cap preparation on each subject, four EEG electrodes (F7, F8, T7, T8) were diverted for use as artifact sensors, by isolating them from the scalp and connecting them to the reference as proposed in previous work (Jorge et al., 2015a).
- **BrainCap MR7Flex:** In this prototype, the EEG leads converged to four PCBs arranged around Cz (Fig. 1-10), which included another 5 kΩ resistor per channel, and then fed two compact flexible ribbon strips (Fig. 1-7,6). As in previous commercial models, four carbon wire loops were attached to the cap to serve as artifact sensors (Warbrick and Schubert, 2020), and exited the cap in a sleeve near Cz (Fig. 1-9,8). Outside the coil, each ribbon strip ended in a compact adapter, to connect to an additional extension cable, and finally to the amplifiers. The distance between the top of the cap (Fig. 1-11) and the amplifier connection was approximately 30 cm. A longer distance of 38 cm was also tested on the phantom, by using longer extension cables to connect to the amplifiers.

**Fig. 1.**
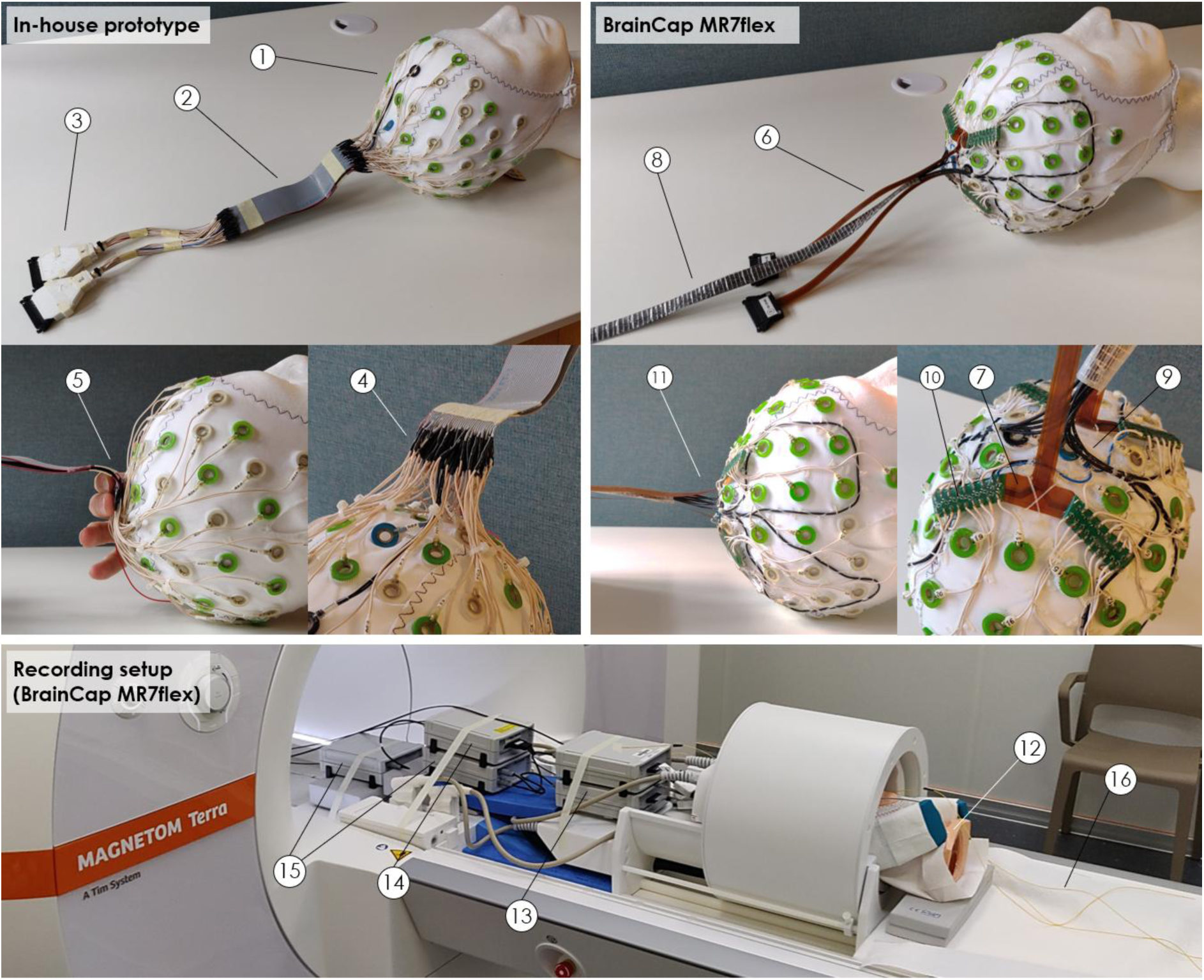
Novel EEG framework explored in this work. Top-left: 64-channel cap adapted in-house from a commercial BrainCap MR model (1), where two flat ribbon cables (each approximately 175 mm long, 42 mm wide, 0.9 mm thick) replace a segment of the leads (2), to fit through the head RF coil. The connectors to the EEG amplifiers (3) used the original printed circuit boards (PCBs) of the base cap, but their housing was replaced by a slimmer 3D-printed version able to fit through the coil gap (when slid open). Each cap lead was soldered to one of the ribbon cable tracks (4), bending outward to the coil gap near the central z-axis (5); the ground and reference channels were positioned at the center of each cable, to reduce induction effects. Four of the original EEG electrodes were diverted for use as artifact sensors (not shown). Top-right: 64-channel BrainCap MR7flex prototype, which uses two compact flexible ribbon strips (each approximately 211 mm long, 10 mm wide, 0.7 mm thick) to pass through the coil gap (6,7), as well as four carbon wire loops distributed over the head (frontal-left, frontal-right, parietal-left, and parietal-right areas) for artifact detection (8,9). The conventional leads stemming from each electrode converge to one of four PCBs arranged around Cz (10), which introduce safety resistors and then feed the two ribbon strips (7), exiting the cap surface near the central z-axis (11). The two ribbon strips (6) and carbon wire sleeve (8) are sufficiently slim to pass through the coil gap in a stack (i.e. on top of each other). Bottom: EEG-fMRI recording setup using the BrainCap MR7Flex, for a test on a head phantom (12). The cap is connected to two 32-channel unipolar amplifiers (13) and one differential amplifier for the carbon loops (14), powered by two batteries (15). Temperature probes were also used in this test (16). The in-house prototype used a similar setup, excluding the differential amplifier and one battery.

Both cap variants connected to two 32-channel BrainAmp MR Plus amplifiers (Brain Products; Fig. 1-13), where the signals were bandpass-filtered (0.016–250 Hz) and digitized (0.5-µV resolution, 5-kHz sampling synchronized with the MR clock). The MR7Flex required an additional BrainAmp ExG MR amplifier to record the carbon loop signals (same filtering, resolution and sampling), powered with a second battery (Fig. 1-14,15). In this compact setup, the cables connecting the cap to the amplifiers were only mechanically fixed at each end (cap and amplifiers), resting on the bottom surface of the narrow coil gap for the larger extent of their length. Past the amplifiers, all signals were optically transmitted. Triggers marking the onset of fMRI volumes were also recorded. Abralyte gel (EasyCap) was used to reduce electrode impedances.

#### 2.1.3. Functional paradigms

Each participant underwent four resting-state acquisitions: 5-min fMRI-only with the 1.6mm protocol (TR=1.05s, 300 volumes) and the 0.8mm protocol (TR=3.52s, 85 volumes), and 8-min EEG-fMRI with each protocol (458 and 136 volumes, respectively). The subjects were asked to lie still with eyes open, and focus on a cross presented on a screen, to minimize movements. The process was repeated for both EEG caps, on separate sessions, and was similarly reproduced for the “reference” participant group (EEG-only, collected outside the scanner room). In the BrainCap MR7Flex EEG-fMRI sessions, a 6.3-min eyes-open/closing paradigm was also run (1.6 mm protocol only), comprising 5 blocks of 32s eyes-open followed by 32s eyes-closed. The transitions were guided by text instructions on the screen, and brief light flashes to signal the end of the eyes-closed periods.

#### 2.1.4. Temperature monitoring

During EEG-fMRI acquisitions, the phantom and first humans were monitored with a 4-channel fiber optic temperature sensor (Neoptix, Québec, Canada). Several points of the EEG acquisition chain were probed across different tests, comprising: (i) EEG electrodes including AF7, Fp2 and FT9, (directly in the gel within the ring), (ii) the ribbon cables (taped to the cable surface), and (iii) the EEG amplifiers (taped to the top surface of the top or bottom amplifier).

### 2.2. Data analysis: safety

The safety evaluation considered the B ^+^, transmit voltage and SAR information logged from each scan, and temperature. From the temperature recordings, average heating rates were estimated for each sequence based on the total temperature variation over the respective acquisition period. Of note, the estimation window was shifted by a manually defined time offset, to account for a delay that was commonly observed between the onset of each acquisition and the start of observable temperature rise.

### 2.3. Data analysis: MRI

Several metrics of data quality and functional sensitivity were estimated for each MRI acquisition, without and with the EEG in place, in specific brain regions and networks. Pre-processing steps were first performed to prepare the fMRI data, and to estimate brain parcellations. In addition to this analysis, the fMRI runs were also studied with ICA and a general linear model (GLM) analysis, to search for the expected networks and responses.

#### 2.3.1. Pre-processing

##### fMRI preparation

Each fMRI run underwent rigid-body motion correction using FSL-FLIRT (v6.0.7.6, Oxford, UK), followed by linear regression to remove slow drift (cosine expansion up to 0.01 Hz) and motion contributions (translations, rotations and their temporal derivatives); the first five volumes were excluded, and the resting-state with-EEG data were cropped to the same duration as the no-EEG (∼5 min), to ensure balanced comparisons. No spatial smoothing was applied at this stage.

##### Parcellations

For each participant, the T_1_-weighted (“MP2RAGE-UNI”) anatomical image was used for cortical and sub-cortical segmentation with Freesurfer (v7.2.0, Charlestown MA, USA). From this segmentation, three parcellations were extracted: (i) a large-scale anatomical parcellation defining the main cortical lobes (occipital, parietal, temporal and frontal), white matter (WM) and cerebellum, built from the Desikan-Killiany (DK) atlas (Desikan et al., 2006), (ii) the functional parcellation proposed by (Yeo et al., 2011) with seven networks (visual, somatomotor, dorsal attention, ventral attention, limbic, frontoparietal and default mode), and (iii) the full DK cortical parcellation with 34 regions per hemisphere (Desikan et al., 2006). Additionally, a “background” region (rectangular cuboid shape) was defined directly on the native space of each image, outside the head and avoiding regions of potential ghosting (adapted for each sequence type, but fixed for all scans of the same sequence).

##### Image registration

For each participant, registration parameters were estimated to allow warping the parcellations from the T_1_ space to the native space of every other image. First, each image type (GRE, B_0_, B_1_, EPIs) was registered between the without- and with-EEG, in-house and MR7Flex conditions using a combination of affine and non-linear symmetric normalization (“SyN”) from ANTs (Avants et al., 2008). Then, in the no-EEG condition where the MP2RAGE was acquired, the T_1_-weighted image was registered to the others using affine boundary-based registration (Greve and Fischl, 2009) in FSL-FLIRT. Specifically for the fMRI runs, the EPI distortions were first estimated using FSL-TOPUP (Andersson et al., 2003), based on the SE-EPI scans. Every registration step was visually inspected and tuned whenever necessary to promote optimal alignments. The brain parcellations were finally warped to the native space of each image by combining the necessary registration steps.

#### 2.3.2. Evaluation of image quality

Image quality was evaluated for the regions of interest (ROI) of the anatomical parcellation. For the B_0_ maps, field heterogeneity was evaluated in terms of the full-width-at-half-maximum (FWHM), i.e. 2√2 *ln*2 multiplied by the standard deviation (STD), of the field values inside each ROI. For the B_1_ ^+^ maps, field strength was evaluated as the ROI-average value, and heterogeneity as the FHWM normalized by the ROI-average value, to decouple the two effects. For the GRE structural, the average magnitude value was computed for each ROI; the background ROI was analyzed following the approach of (Aja-Fernandez et al., 2008), which accommodates the presence of both residual signal and noise in the ROI (i.e. a full Rician distribution (Cárdenas-Blanco et al., 2008)), and estimates the noise STD based on local sample metrics (local mean, STD and RMS) in a sliding window approach; the spatial signal-to-noise ratio (SNR) was then computed as the ratio between the average value in each ROI and the noise STD of the background ROI. For the fMRI runs, the average value, background noise STD and spatial SNR were similarly computed for each timepoint, and then averaged across time; temporal SNR was estimated for each voxel of the WM region as the temporal mean divided by the temporal STD, and then averaged across voxels.

#### 2.3.3. Evaluation of functional sensitivity

Functional sensitivity was primarily evaluated for the ROIs (networks) of the functional parcellation, on the pre-processed fMRI runs. Two metrics were estimated: (i) the fractional amplitude of low frequency fluctuations (fALFF), estimated for each voxel as the ratio between the sum of spectral amplitudes in 0.01–0.08 Hz and the sum over 0–0.25 Hz (Zou et al., 2008), and (ii) intra-network consistency (INC), adapted from the concept of ROI consistency (Korhonen et al., 2017), and estimated for each ROI (Yeo network) as the average Pearson correlation between each voxel timecourse and the ROI-average timecourse, after temporal bandpass filtering the data at 0.01–0.08 Hz. Considering the nature of these metrics (Korhonen et al., 2017; Zou et al., 2008), it was hypothesized that a loss in functional sensitivity due to the presence of EEG should lead to a decrease in fALFF across the brain, particularly the cortex, and to a decrease in INC within each ROI, since each ROI corresponds to a meaningful functional network with correlated activity. Additionally, on the DK parcellation, we conducted a pair-wise connectivity analysis following the well-established procedure previously adopted in (Wirsich et al., 2021): the fMRI data were motion- and slice timing-corrected, and the first 5 volumes were excluded; confounds from slow drift, head motion, average WM, average CSF and average GM signals were regressed out (based on WM, CSF and GM ROIs defined from the Freesurfer segmentation), and the data were bandpass-filtered (0.009–0.08 Hz); pair-wise connectivity was then estimated in terms of the Pearson correlation between the average timecourse of each DK region.

In parallel, the resting-state fMRI data was also studied with ICA, and the eyes-open/closing data was studied with both ICA and a GLM analysis (Worsley and Friston, 1995). For ICA, in-between the motion correction and nuisance regression steps of the pre-processing, the data were slice timing-corrected and spatially smoothed (Gaussian, 1.6 mm FWHM). Each run was then decomposed with spatial ICA (extended Infomax algorithm (Lee et al., 1999)), after dimensionality reduction to 20 components. The resting-state decompositions were inspected to identify two typical networks: default mode (DMN), and a visual network (primary, secondary, or a mix) (Smith et al., 2009); the eyes-open/closing data were instead inspected to identify ICs whose timecourse was synchronized with the paradigm.

For the GLM analysis, the eyes-open/closing fMRI runs were motion- and slice timing-corrected, spatially smoothed (Gaussian, 1.6 mm FWHM), and the first 5 timepoints were excluded. The base GLM model included slow drift and motion contributions (similar to those described in 2.3.1). For hemodynamic activity, three contributions were introduced: (i) response to visual input change upon closing the eyes, modeled as a train of delta functions (“sticks”) marking the timepoints of eyes closing, convolved with a single-gamma hemodynamic response function (HRF) and its derivative (Jezzard et al., 2001); (ii) response to visual input change upon opening the eyes, modeled in a similar way; and (iii) fluctuations associated with alpha activity, modeled by extracting the alpha power timecourse of the EEG IC with strongest alpha modulations, convolving it with the HRF and its derivative, and orthogonalizing with respect to (i) and (ii). After GLM fitting, a Z-score activation map was estimated for the alpha regressor. Moreover, to assess what proportion of the fMRI information could be meaningfully described by alpha, the explained variance of this regressor was calculated based on the adjusted coefficient of determination (R^2^) of the model fit with and without the regressor (Jorge et al., 2013). For comparison against a “control” equivalent, the same procedure was then repeated with a randomized version of the alpha regressor, obtained by preserving its spectral magnitude while randomizing the spectral phase (repeated for 100 randomization trials per recording).

### 2.4. Data analysis: EEG

To evaluate the impact of the MRI environment on the EEG data, before and after artifact reduction, different correction steps were sequentially applied, while evaluating several properties of the EEG data with respect to the “reference” participant group collected outside the scanner room.

#### 2.4.1. Artifact reduction approach

The EEG data were processed with several techniques derived from the existing literature, including our previous works. The choice of methods and parameters was guided by visual inspection of the EEG spectra, benchmarked against phantom data, inside-scanner without-scanning human data, and outside-scanner data (e.g. Supp. Fig. 5), seeking to maximize the removal of artifactual content while preserving real EEG content. To avoid biases, no comparisons to the reference group were made while correcting the EEG-fMRI group. The procedure (detailed in Supp. Table II) included the following steps, in order: GA and PA trigger estimation; GA correction based on average artifact subtraction (Allen et al., 2000) followed by optimal basis set removal (Niazy et al., 2005); PA correction using a k-means clustering-based approach (Gonçalves et al., 2007; Jorge et al., 2019); downsampling to 200 Hz; MA correction (including other artifact residuals) by adaptive regression of the artifact sensor signals of each cap (Jorge et al., 2015a; Masterton et al., 2007); bandpass filtering to 0.75–70 Hz (chosen as the range of interest for the quality analysis), bad channel interpolation and re-referencing to the channel average; ICA-based correction to exclude components associated to artifact residuals, eye and muscle activity.

Accordingly, the reference group underwent only temporal downsampling (200 Hz), bandpass filtering (0.75–70 Hz), bad channel interpolation, average channel re-referencing, and then ICA to exclude spurious contributions from e.g. eye and muscle activity.

#### 2.4.2. Evaluation of signal quality and characteristics

The EEG evaluation metrics covered spectral, spatial and temporal properties. For the spectral metrics, a magnitude spectrum was first estimated for each channel using Welch’s method (10 s Hann window, 50% overlap); the average magnitude in different frequency windows was then extracted, including the delta band (0.75–4 Hz), theta (4–7 Hz), alpha (7–13 Hz), beta (13–30 Hz), gamma (30– 70 Hz), and the full retained bandwidth (0.75–70 Hz).

The spatial and temporal properties were analyzed in the framework of a microstate analysis in the 1–40 Hz band; microstates provide a well-studied summary description of whole-scalp EEG recordings, based on their dominant topographies and respective temporal dynamics (Michel and Koenig, 2018; Pascual-Marqui et al., 1995). To this end, after each artifact correction step, topographies were estimated based on k-means clustering of the GFP peaks of the participant group (all recordings bandpass-filtered at 1–40 Hz, GFP-normalized and then concatenated in time), using van Vliet’s module for MNE-Python (Poulsen et al., 2018; van Vliet, 2022). The clustering analysis was fixed at 4 classes, for easier referencing with respect to the majority of the existing literature (Michel and Koenig, 2018), to promote more stable clustering estimations, particularly on artifact-disrupted signals, and for higher simplicity in terms of organization and interpretation of the results. Once estimated, the topographies were back-fitted to the EEG of each subject to estimate each microstate’s duration (average duration that the state remains stable), occurrence (number of times per second that the state reappears) and coverage (fraction of total time dominated by the state) (Michel and Koenig, 2018).

The presence of alpha wave modulation (Adrian and Matthews, 1934) was studied on the EEG of the eyes-open/closing runs. During the ICA-based correction step, the alpha power timecourse of each IC was computed, to investigate the presence of ICs showing alpha synchrony with the paradigm.

### 2.5. Data analysis: statistical significance

Given the relatively moderate group sizes, the report of results below favored showing all individual data points whenever possible, and drawing global observations accordingly. These observations were complemented with more quantitative summary metrics, namely average values across participants, and the significance of group-level differences was assessed via t-tests (significance threshold: *p* < 0.05).

## 3. Results

### 3.1. Safety

All MRI sequences that were intended for use with EEG exhibited a B ^+^ below 1 µT, with the fMRI sequences at approximately 0.6 µT (Table I). On average across subjects, the presence of the EEG prompted an increase in transmit voltage of approximately 10% by the scanner calibration procedures (*p* < 0.01, similar in both caps), accompanied by an increase in SAR of 22–27% across sequences (Table I) – which nonetheless remained within the limits of the scanner’s “normal operation mode” (20 W/Kg at 10 s and 10 W/Kg at 6 min, with the fMRI sequences only reaching ∼3.8 W/Kg).

**Table I.** B ^+^, transmit voltage and SAR for the MRI sequences used in this study.

| Sequence | $B_1^+$<sub>RMS</sub><br>(μT) | Transmit voltage (V) | | SAR (W/Kg) | |
| --- | --- | --- | --- | --- | --- |
|  |  | Reference | Adjusted | Reference | Adjusted |
| GE-EPI 1.6 mm | 0.62 | 221 – 248 | 251 – 271 | 2.45 – 3.05 | 3.20 – 3.73 |
| GE-EPI 0.8 mm | 0.60 |  |  | 2.45 – 3.04 | 3.25 – 3.79 |
| T<sub>1</sub> (MP2RAGE) | 1.00 |  |  | 6.42 – 7.89 | Not tested |
| GRE structural | 0.27 |  |  | 0.46 – 0.58 | 0.60 – 0.70 |
| B<sub>1</sub> (SA2RAGE) | 0.41 |  |  | 1.07 – 1.36 | 1.39 – 1.62 |

**Note:** The values reported for $B_1^+$ <sub>RMS</sub> were obtained in a preliminary phantom session. The value ranges reported for the transmit voltage and SAR cover all human participants and both caps. The “reference” case corresponds to $V_{\text{ref}}$ , and the “adjusted” case to $V_{\text{adj}}$ (see Methods section 2.1.1). The $B_1^+$ <sub>RMS</sub> and SAR values reported correspond to the maximum observed between two measures provided by the MRI system, corresponding to averaging timescales of 10 s and 6 min, respectively. The SAR limits imposed by the system during the tests were the standard 20 W/Kg at 10 s and 10 W/Kg at 6 min.

Overall, the temperature tests did not indicate safety risks for the subjects (Table II). No substantial heating effects were observed on any of the electrodes tested, with heating rates that did not exceed 0.05°C/min. Consistent with this, none of the participants reported any heating sensations on the scalp during the recordings, despite being instructed to remain attentive to this possibility. On the other hand, the EEG amplifier surfaces did show stronger heating with the BrainCap MR7Flex when using shorter extension cables (detailed in section 2.1.2), reaching up to 0.37°C/min during GE-EPI. However, the same setup with longer extensions had substantially lower heating – below 0.1°C/min, similar to the in-house cap. On the flexible ribbon strips (MR7Flex), the heating rates were between 0.06 and 0.10°C/min during GE-EPI; interestingly, a phantom test with SE-EPI elicited visibly stronger heating there, up to 0.24°C/min (Table II). Despite its considerably higher resolution, the 0.8 mm fMRI protocol ran at a similar SAR to the 1.6 mm protocol, and was not found to cause stronger heating effects overall.

**Table II.**
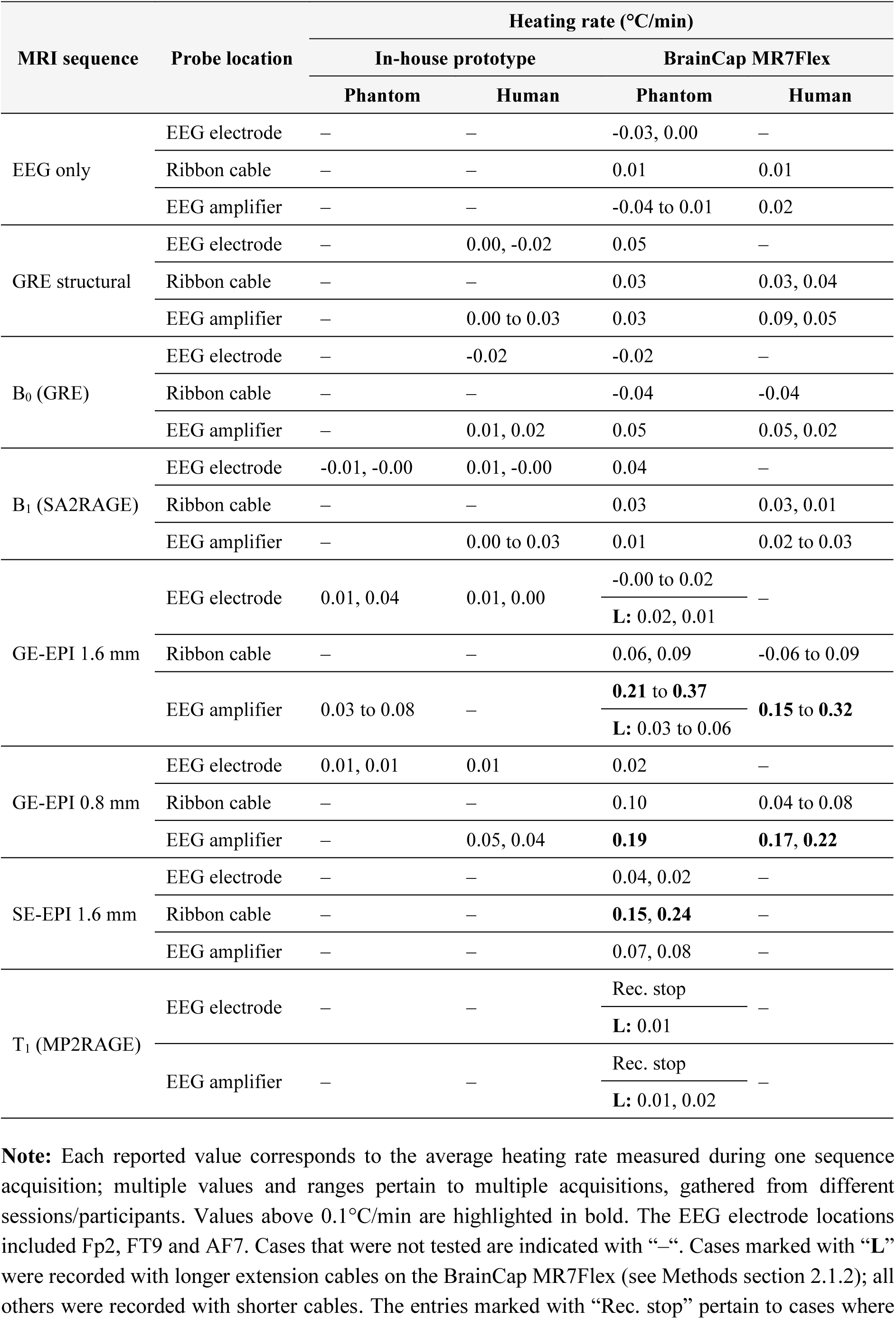
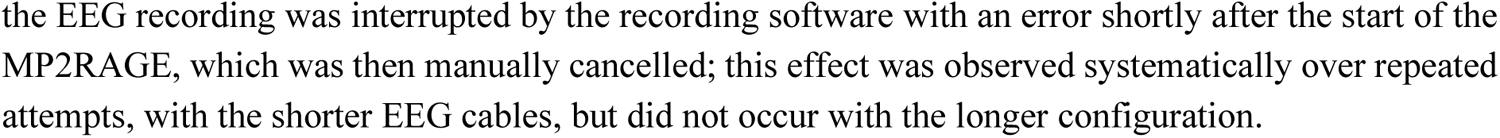
Heating effects measured during EEG and concurrent (f)MRI acquisitions.

| MRI sequence | Probe location | Heating rate (°C/min) |  |  |  |
| --- | --- | --- | --- | --- | --- |
|  |  | In-house prototype |  | BrainCap MR7Flex |  |
|  |  | Phantom | Human | Phantom | Human |
| EEG only | EEG electrode | – | – | -0.03, 0.00 | – |
|  | Ribbon cable | – | – | 0.01 | 0.01 |
|  | EEG amplifier | – | – | -0.04 to 0.01 | 0.02 |
| GRE structural | EEG electrode | – | 0.00, -0.02 | 0.05 | – |
|  | Ribbon cable | – | – | 0.03 | 0.03, 0.04 |
|  | EEG amplifier | – | 0.00 to 0.03 | 0.03 | 0.09, 0.05 |
| B<sub>0</sub> (GRE) | EEG electrode | – | -0.02 | -0.02 | – |
|  | Ribbon cable | – | – | -0.04 | -0.04 |
|  | EEG amplifier | – | 0.01, 0.02 | 0.05 | 0.05, 0.02 |
| B<sub>1</sub> (SA2RAGE) | EEG electrode | -0.01, -0.00 | 0.01, -0.00 | 0.04 | – |
|  | Ribbon cable | – | – | 0.03 | 0.03, 0.01 |
|  | EEG amplifier | – | 0.00 to 0.03 | 0.01 | 0.02 to 0.03 |
| GE-EPI 1.6 mm | EEG electrode | 0.01, 0.04 | 0.01, 0.00 | -0.00 to 0.02<br>**L: 0.02, 0.01** | – |
|  | Ribbon cable | – | – | 0.06, 0.09 | -0.06 to 0.09 |
|  | EEG amplifier | 0.03 to 0.08 | – | **0.21 to 0.37**<br>**L: 0.03 to 0.06** | **0.15 to 0.32** |
| GE-EPI 0.8 mm | EEG electrode | 0.01, 0.01 | 0.01 | 0.02 | – |
|  | Ribbon cable | – | – | 0.10 | 0.04 to 0.08 |
|  | EEG amplifier | – | 0.05, 0.04 | **0.19** | **0.17, 0.22** |
| SE-EPI 1.6 mm | EEG electrode | – | – | 0.04, 0.02 | – |
|  | Ribbon cable | – | – | **0.15, 0.24** | – |
|  | EEG amplifier | – | – | 0.07, 0.08 | – |
| T<sub>1</sub> (MP2RAGE) | EEG electrode | – | – | Rec. stop<br>**L: 0.01** | – |
|  | EEG amplifier | – | – | Rec. stop<br>**L: 0.01, 0.02** | – |

**Note:** Each reported value corresponds to the average heating rate measured during one sequence acquisition; multiple values and ranges pertain to multiple acquisitions, gathered from different sessions/participants. Values above 0.1°C/min are highlighted in bold. The EEG electrode locations included Fp2, FT9 and AF7. Cases that were not tested are indicated with “–”. Cases marked with “**L**” were recorded with longer extension cables on the BrainCap MR7Flex (see Methods section 2.1.2); all others were recorded with shorter cables. The entries marked with “Rec. stop” pertain to cases where
the EEG recording was interrupted by the recording software with an error shortly after the start of the MP2RAGE, which was then manually cancelled; this effect was observed systematically over repeated attempts, with the shorter EEG cables, but did not occur with the longer configuration.

### 3.2. Usability

The EEG-fMRI participant group included several individuals that had never undertaken an MRI acquisition, and only two had previously undergone simultaneous EEG-fMRI. The recordings were nonetheless well tolerated by all participants, despite the presence of not only the EEG cap but also the respiratory belt and finger photoplethysmography sensor. The EEG gel preparation was not found more challenging with the new cap prototypes than with a standard BrainCap MR. The process of setting up the participant in the head RF coil and handling the cables within this narrow space did present new practical difficulties compared to more spacious coils, but did not block the procedure in any of the participants. Additional practical difficulties were found in cap cleaning (gel removal), as we avoided submerging the soldered connections of the in-house prototype (Fig. 1-4) or the PCBs of the MR7Flex (Fig. 1-10), respectively. Similarly, this lengthened the task but did not prevent its completion.

### 3.3. MRI data quality

The volume occupied by the EEG cap and leads introduced a small but measurable constraint for head placement in the RF coil, requiring on average across participants a displacement of the head in the head-to-feet direction of 2.4 cm for the in-house prototype (1.7 cm for the MR7Flex), and of 0.6 cm (respectively 0.3 cm) in the posterior-to-anterior direction, as well as a sagittal rotation of 14.3° (respectively 11.2°). This physical constraint did not prevent any image acquisitions, but did influence the distortions in lower head areas, motivating the use of non-linear methods for registration between the images with and without EEG, as described in section 2.3.1.

Upon visual inspection, the quality of the MRI data, without EEG, was in line with previous datasets acquired by our group at 7T with a 32-channel RF array, and superior to data obtained with an 8-channel array (Jorge et al., 2015b). The introduction of the EEG setup did cause visible signal reductions, particularly in superior-parietal regions, with both caps (Fig. 2, Supp. Fig. 1). Similar to the observations of (Meyer et al., 2020), these reductions were relatively mild, and without localized accentuated drops such as observed in previous setups (Jorge et al., 2015b; Mullinger et al., 2008b). The B_1_ ^+^ maps revealed reductions in the same areas; the B_0_ maps showed increased heterogeneity at the head surface (skin, etc.), possibly close to EEG electrodes and gel, but these perturbations did not appear to reach inside the brain (Fig. 2, Supp. Fig. 1). For some participants, the 0.8 mm fMRI data showed important ghosting artifacts along the anterior-posterior direction in lower slices, but occurring both with and without EEG.

**Fig. 2.**
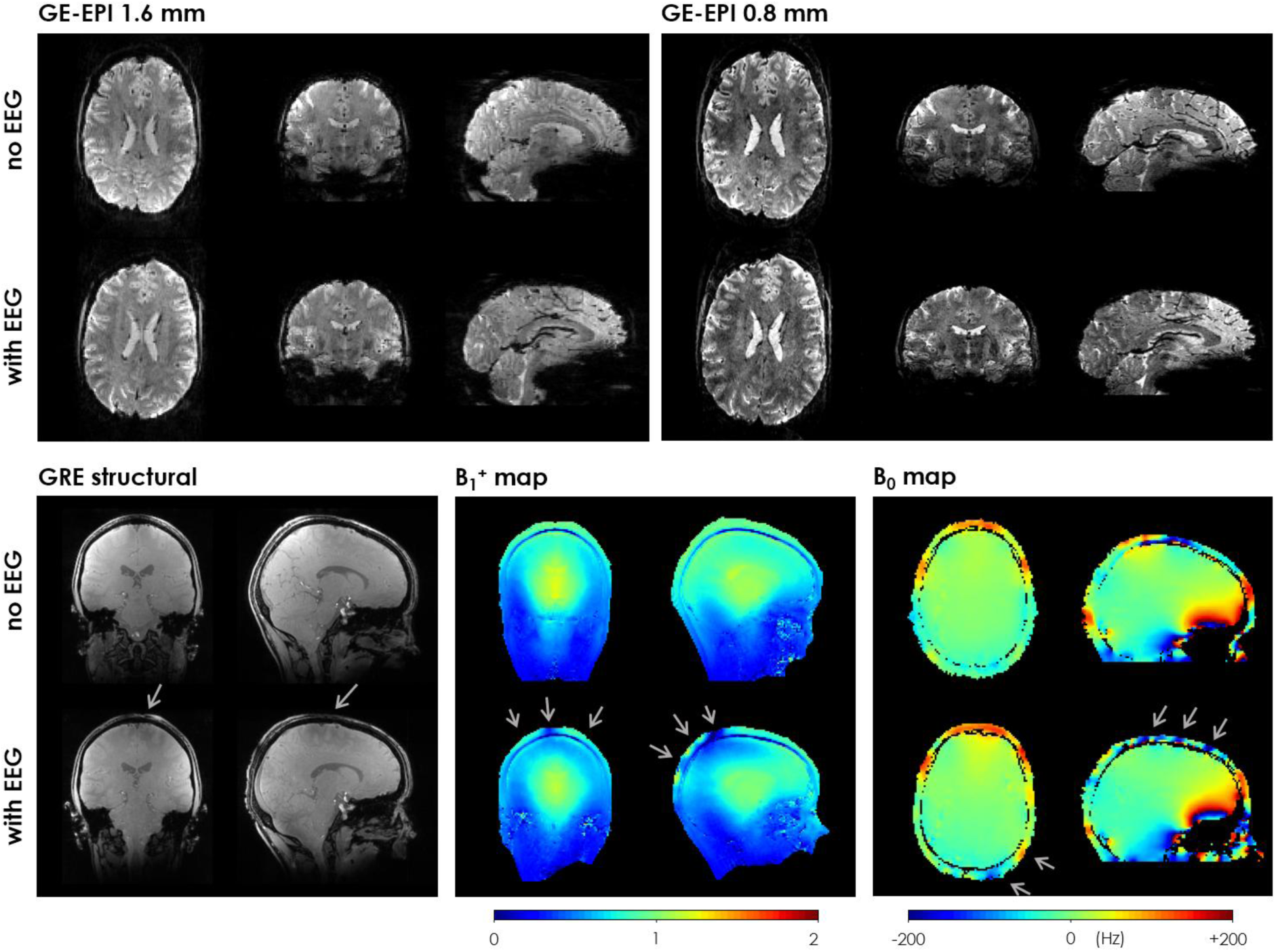
MRI data obtained from an example subject with and without the EEG system in place. This example used the in-house EEG cap prototype. The slices shown include some of the most relevant differences found between the with- and without-EEG conditions. Top: GE-EPI volumes from the fMRI acquisitions (left: 1.6mm isotropic resolution; right: 0.8 mm isotropic resolution). Bottom: GRE-based anatomical image and field maps. The B_1_^+^ map is expressed as a fraction of the nominal flip angle. Both maps were masked to exclude background voxels. The grey arrows point to interference effects likely caused by the presence of the EEG system.

The quantitative analysis was generally consistent with the visual observations, although with less accentuated variations at ROI level than seen on the example slices of Fig. 2 and Supp. Fig. 1, which were selected specifically to illustrate the most relevant alterations. Across the brain on average, the B_0_ field inhomogeneity showed a tendency for increasing with the introduction of the EEG, by approximately 7.5% for the in-house prototype and 4.3% for the BrainCap MR7Flex, although non-statistically significant with high variability across subjects (Fig. 3, 1^st^ row). The B_1_ ^+^ strength had consistent decreases with EEG for every participant, brain region and both caps (Fig. 3, 2^nd^ row): when keeping the same transmit voltage (V_ref_), B_1_ ^+^ strength decreased by 22.7% with the in-house cap and 17.6% with the MR7Flex, across the brain on average; when adjusting to V_adj_, the reduction was systematically (though not fully) mitigated, to 14.9% and 8.6%, respectively (all statistically significant reductions, *p* < 0.01). The most affected regions were the occipital and parietal lobes and the cerebellum. B_1_ ^+^ heterogeneity also tended to increase, for the whole brain (*p* < 0.02 in both caps) and particularly in the parietal lobe with the in-house cap, but with substantial variability across regions and subjects (Fig. 3, 3^rd^ row). In the GRE structural data, the EEG caused significant signal losses in all subjects and brain regions, averaging at 28.4% with the in-house cap and 26.8% with the MR7Flex, when fixed at V_ref_, and only slightly mitigated to 25.2% and 23.4% with V_adj_ (*p* < 0.01 in all cases; Fig. 3, 4^th^ row). The background noise STD was also reduced, yet only by approximately 6 to 10% across cases (*p* < 0.05; Fig. 3, 4^th^ row-right). Hence, the image SNR experienced reductions of 20.0% with the in-house cap and 22.0% with the MR7Flex, at V_ref_, and slightly improved to 15.1% and 17.6% at V_adj_ (*p* < 0.02; Fig. 3, 5^th^ row). The SNR loss was highest in the parietal lobe (∼24%), but this region also had a high SNR to begin with.

**Fig. 3.**
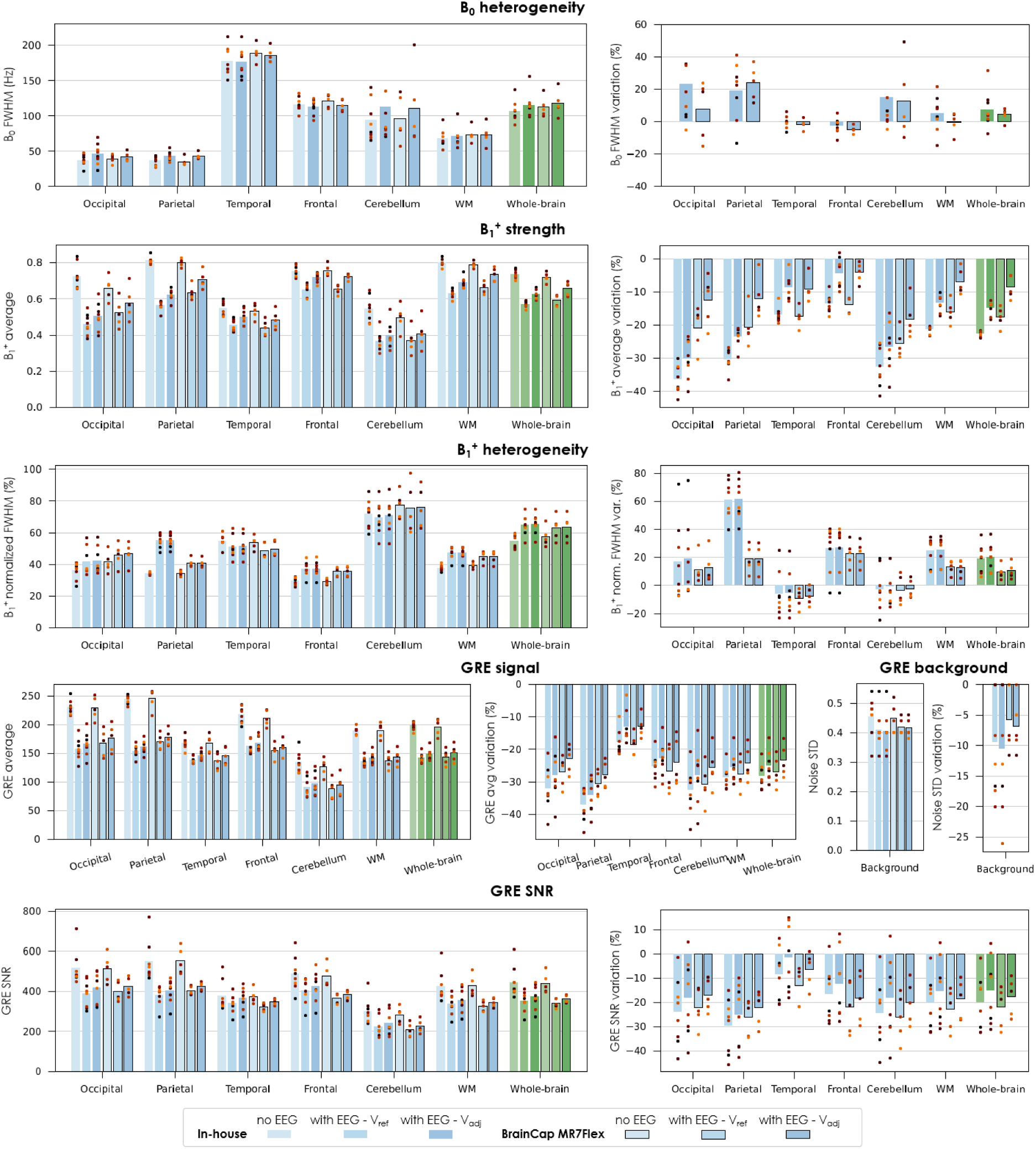
Impact of EEG on MRI data quality for different brain regions, evaluated on B_0_ mapping, B_1_^+^ mapping and GRE-based structural data. The images were acquired without and with EEG, including each of the two tested caps. The B_1_^+^ and GRE images with EEG were acquired twice, with different transmit voltage (V_ref_ and V_adj_ – see Methods section 2.1.1). The B_1_^+^ strength is expressed as a fraction of the nominal flip angle. B_0_ and B_1_^+^ heterogeneity was evaluated in terms of the FWHM of the map’s value distribution within each region; in the case of the B_1_^+^, this was then divided by the average value, to separate the concurrent effect of average B_1_^+^ reduction. The GRE SNR was estimated by dividing the average intensity in each region by the noise STD of the background region. For each of the quality metrics, the relative variation upon introducing the EEG system (with respect to no-EEG) is also plotted on the side. Each bar represents the average across subjects; the dot markers represent the individual subjects.

Regarding the GE-EPI data, overall, the 1.6 mm data had higher spatial and temporal SNR than the 0.8 mm. For the 1.6 mm acquisitions, the signal losses due to EEG averaged at 26.6% with the in-house cap and 24.6% with the MR7Flex, and the background noise STD reductions at 11.2% and 8.6% (Fig. 4, 1^st^ row), resulting in spatial SNR losses of 17.0% and 16.7% (*p* < 0.03 for both; Fig. 4, 2^nd^ row). The temporal SNR in WM experienced smaller average reductions, with high variability across subjects: 10.7% (*p* < 0.01) and 6.2% (non-significant). The 0.8 mm data exhibited similar losses in signal amplitude, but the background noise STD was less reduced (3.1% and 0.1%; Fig. 4, 3^rd^ row). As a result, the spatial SNR was more penalized, by 25.2% and 25.8% (*p* < 0.01 in both; Fig. 4, 4^th^ row). The averages losses in temporal SNR were also stronger than at 1.6 mm, yet not to the same extent: 16.9% (*p* < 0.01) and 10.6% (non-significant).

**Fig. 4.**
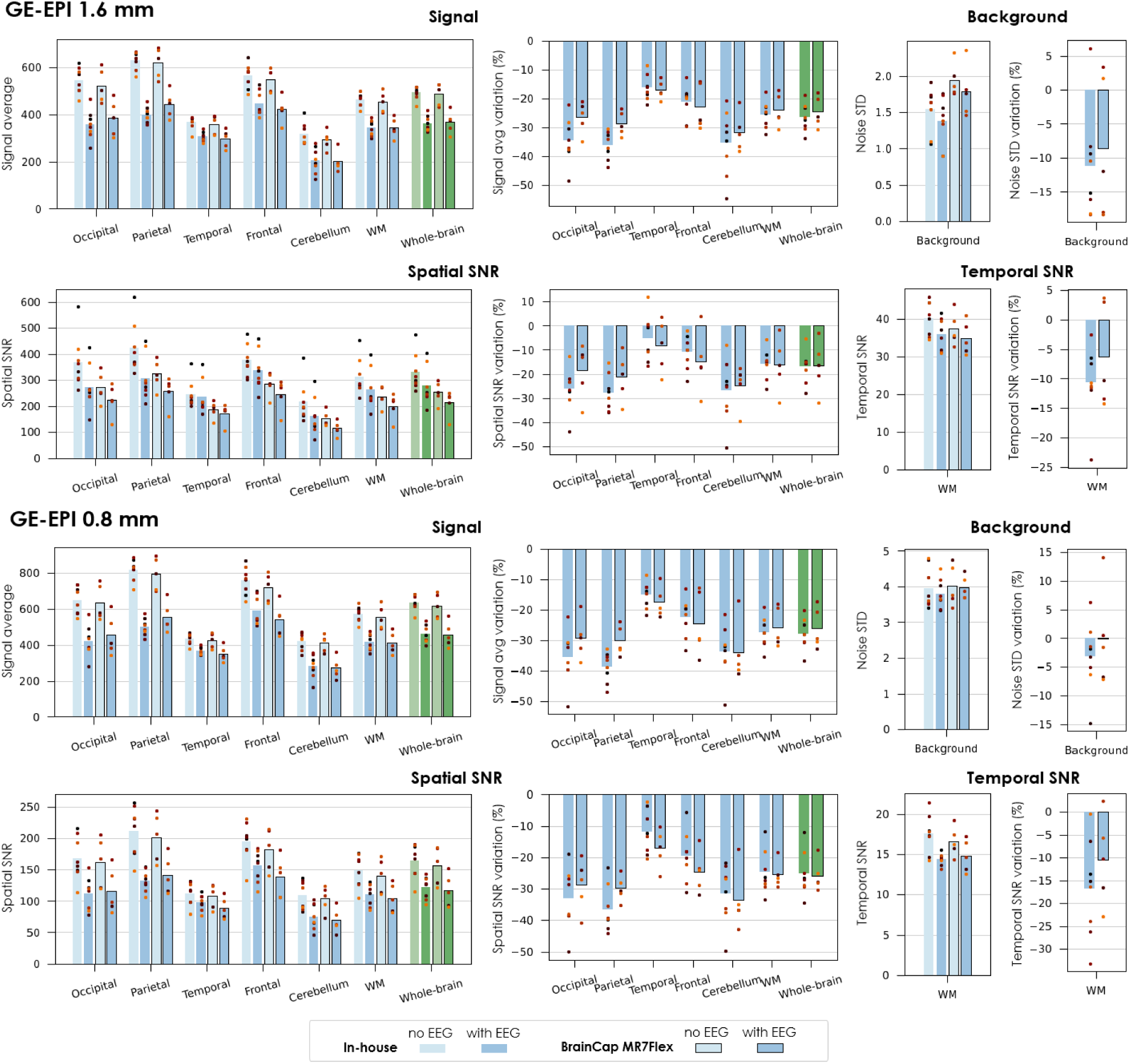
Impact of EEG on fMRI data quality for different brain regions, evaluated on the 1.6 mm and 0.8 mm GE-EPI acquisitions. The images were acquired without and with EEG, including each of the two tested caps. The transmit voltage was calibrated for each case (V_ref_ without EEG, V_adj_ with EEG). The GE-EPI signal average was computed across time and voxels of each region; the STD was computed across voxels of the background and then averaged across time. The spatial SNR was estimated by dividing the signal average in each brain region by the noise STD of the background region, at each timepoint, and then averaging across time. The temporal SNR was estimated for each voxel as the mean divided by the STD across time, and then averaged across voxels. For each of the quality metrics, the relative variation upon introducing the EEG system (with respect to no-EEG) is also plotted on the side. Each bar represents the average across subjects; the dot markers represent the individual subjects.

The functional sensitivity metrics behaved as expected in several aspects: in the 1.6 mm data, within the defined networks, the fALFF range (approximately 0.30–0.36) was in line with previous fMRI data at 7T (Bréchet et al., 2019), whereas it was lower in the background ROI, as would be expected for a region dominated by random noise (Zou et al., 2008) (Fig. 5, 1^st^ row). The INC, estimated at 0.21–0.29 for the network ROIs, was also in line with previous reports on functionally-defined ROIs such as the Brainnetome atlas (Korhonen et al., 2017); as expected, it was lower for the whole-brain ROI (as it contains multiple non-covarying networks), and substantially lower in the background (Fig. 5, 2^nd^ row). The 0.8 mm protocol achieved lower fALFF and INC values in general, but with similar behavior (Fig. 5, 4^th^, 5^th^ rows). Overall, the variability across subjects and sessions appeared dominant with respect to any potential systematic variations due to EEG, which were non-statistically significant (except for the fALFF in some networks with the MR7Flex, 0.8 mm). The connectivity analysis performed on the DK cortical parcellation yielded group connectivity matrices that, overall, exhibited similar dominant patterns in either fMRI protocol, with and without EEG, with Pearson correlation values ranging from −0.63 to +0.93 (Supp. Fig. 2 and Supp. Fig. 3, top row). The group-average matrices without vs. with EEG showed a spatial correlation of 0.94 (1.6 mm protocol) and 0.91 (0.8 mm protocol), which are on the upper end of previous reports across field strengths (Wirsich et al., 2021). The differences in pair-wise connectivity between the with-EEG and no-EEG conditions were generally smaller than the respective STD across subjects (Supp. Fig. 2 and Supp. Fig. 3, middle row); in both fMRI protocols, several region pairs showed significant differences (*p* < 0.05), yet none remained significant after adjusting for multiple comparisons (false discovery rate), except for one pair in the 0.8 mm data (Supp. Fig. 2 and Supp. Fig. 3, bottom row). An additional check on the average absolute connectivity change per DK region indicated that no particular regions were visibly more strongly affected by the EEG than others, in either fMRI protocol (Supp. Fig. 4). Finally, the ICA investigation allowed a robust recovery of components matching the spatial configuration of the DMN and visual networks, in all subjects (except one in the 0.8 mm data) (Fig. 5, 3^rd^, 6^th^ rows); the respective maps did not exhibit visible systematic deviations between the no-EEG and with-EEG conditions.

**Fig. 5.**
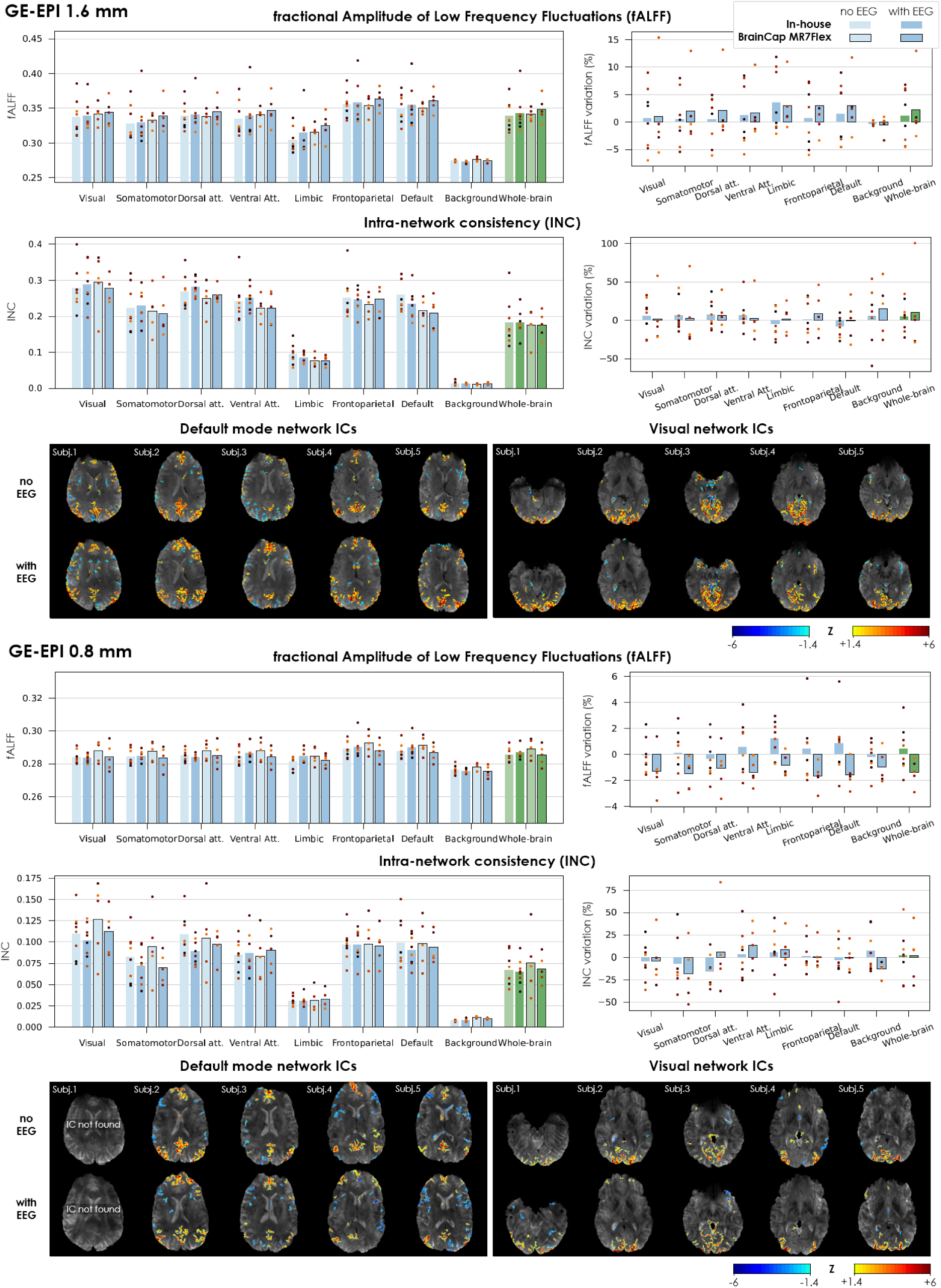
Impact of EEG on fMRI sensitivity for different brain networks, evaluated on the 1.6 mm and 0.8 mm GE-EPI acquisitions. The images were acquired without and with EEG, including each of the two tested caps. The transmit voltage was calibrated for each case (V_ref_ without EEG, V_adj_ with EEG). All metrics were extracted after fMRI pre-processing (motion correction, spatial smoothing, slice-timing correction, temporal cropping, and drift + motion regression). For each of the sensitivity metrics, the relative variation upon introducing the EEG system (with respect to no-EEG) is also plotted on the side. Each bar represents the average across subjects; the dot markers represent the individual subjects. The ICA maps shown are the most representative of the DMN and visual networks (including sub-regions of the primary and/or secondary visual cortex) found for five example subjects without and with EEG (BrainCap MR7Flex). All IC maps were converted to Z-score maps and thresholded at |*Z*| > 1.4.

**Fig. 6.**
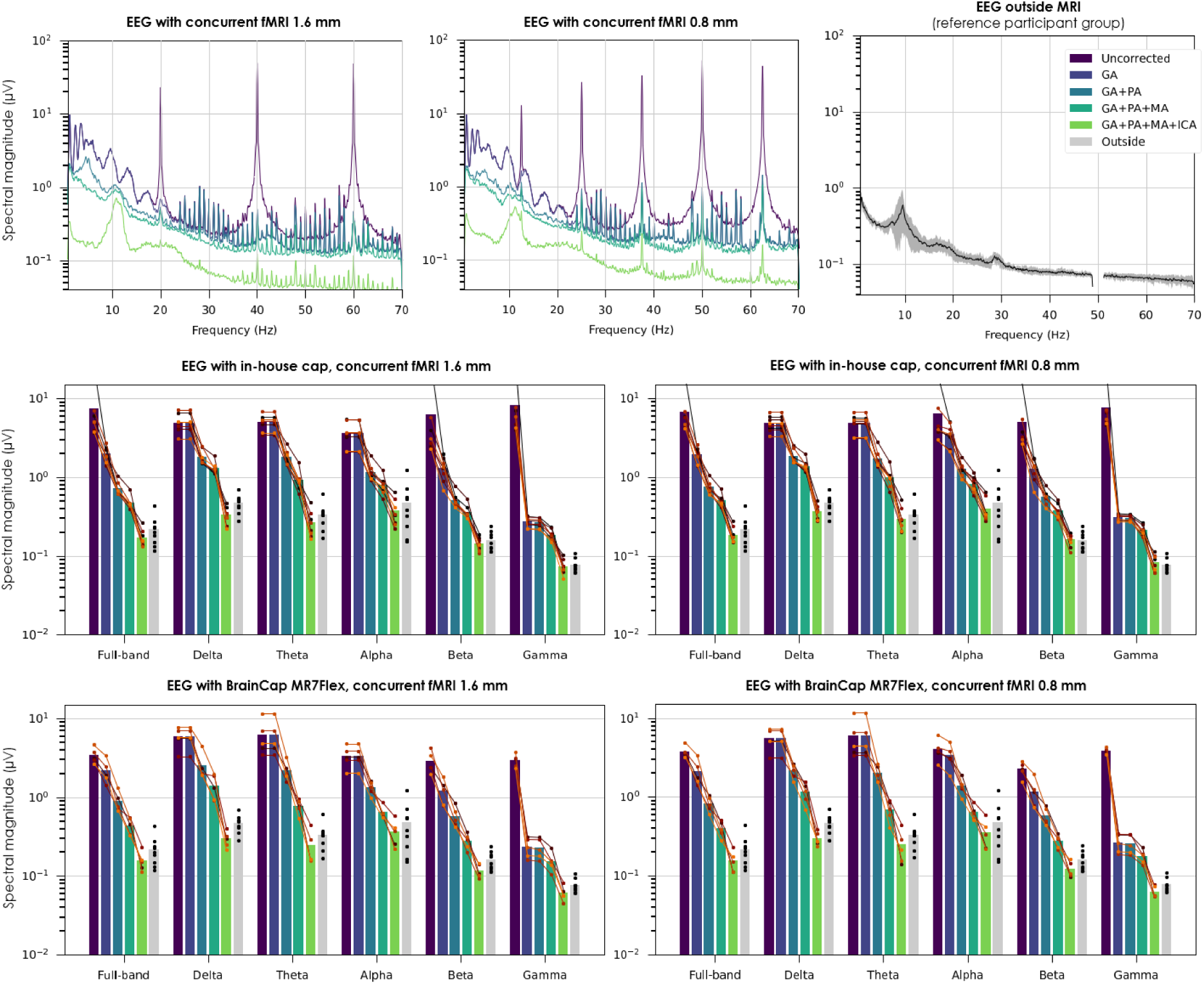
Impact of MRI-related artifacts and the respective correction steps on EEG data quality, in terms of spectral content. First row, left and middle: signal spectra in an example participant at rest (in-house EEG cap), with either fMRI protocol; for consistency across correction steps, all cases were first downsampled to 200 Hz, bandpass-filtered to 0.75–70 Hz and re-referenced to the channel average; magnitude spectra were then estimated using Welch’s method (10 s Hann window, 50% overlap) for each channel, and averaged across channels. For reference, the slice GA peaks are expected at 20 Hz for the 1.6 mm fMRI protocol (respectively 12.5 Hz for the 0.8 mm protocol) and harmonics; the volume GA peak, if present, is expected at 0.95 Hz (respectively 0.28 Hz). First row, right: signal spectrum for the reference participant group recorded outside MRI (non-shielded room), pre-processed in a similar manner, except for an additional 50 Hz notch filter; the black line denotes the average across channels and participants; the gray margin denotes the STD across participants. Second and third row: impact of artifact correction on different frequency bands across all subjects, for each fMRI acquisition protocol and EEG cap prototype. The spectral magnitudes were computed through the same Welch-based approach as the spectra, followed by averaging within the frequency bands of interest. Each bar represents the average across subjects; the colored dot markers and lines represent the individual subjects. The reference group is also shown, in gray, for comparison.

### 3.4. EEG data quality

All EEG data recorded with concurrent fMRI (phantom and human, both caps, resting-state and eyes open/closing runs) were checked for the presence of amplitude saturation. Of 35 recordings, saturation instances were detected in 5, of which 4 used the 0.8 mm protocol. In the majority of cases, this affected no more than 1 channel per recording – not always the same, but typically coinciding with a contact impedance close to or above 100 kΩ. When present, the saturation periods tended to occur on the largest peak of the slice GA waveform (linked to the slice selection block), and typically affected less than 2% of the time samples of the channel.

Phantom recordings in-scanner, without concurrent MRI, exhibited numerous peaks over a wide range of frequencies above ∼10 Hz (Supp. Fig. 5, top-right), consistent with Helium coldhead contributions previously observed at 7T (Jorge et al., 2015b; Mullinger et al., 2008a). The fMRI scans then introduced prominent peaks (GAs) matching the inverse of the slice TR and harmonics. The human recordings exhibited the same contributions, as well as PAs comprising a peak at the fundamental cardiac frequency and harmonics, with wider inflections spanning up to ∼30 Hz (Supp. Fig. 5, bottom), completely superseding alpha and beta-wave contributions seen outside the scanner (Supp. Fig. 5, bottom-right). These observations were consistent with the spectral profiles observed across correction steps, where the alpha peak only started becoming visible after GA+PA correction, and beta was only discernible after the full correction (Fig. 6, 1^st^ row; Supp. Fig. 6). The peaks associated to EAs could be partially reduced by the reference sensor-based MA correction, and then further (though not fully) by ICA (Supp. Fig. 6). These effects were also reflected on the different frequency bands of interest (Fig. 6, 2nd and 3rd rows). All correction steps played a substantial role in reducing spectral magnitude, with varying impact across frequency bands depending on the characteristics of the respective artifacts. Encouragingly, at the end of the correction pipeline, the band-specific power values of the group became very comparable to the reference group, with a statistically significant difference (*p* < 0.05) only found in the delta band.

In the microstate analysis, the reference participant group yielded topographies that closely matched the four “canonical states” A–D typically found at rest (Michel and Koenig, 2018) (Fig. 7, top-middle). The EEG data with concurrent fMRI did yield topographies resembling microstates A–C at all correction steps; however, the topography assigned to D was dominated by a left-to-right distribution typically linked to induction artifacts (Jorge et al., 2015a; Yan et al., 2010), that only gave place to the more characteristic microstate D after full correction (Fig. 7, top-left). The recovery of all four canonical topographies was consistently successful for both caps and both fMRI protocols after full artifact correction (Fig. 7, top-right). The temporal properties of the microstates showed substantial variability across subjects, in both reference and EEG-fMRI groups (Fig. 7, bottom). Nonetheless, across correction steps, the average profiles of the EEG-fMRI runs became increasingly closer to the reference, with certain metrics undergoing changes of 20–30% in several microstates, comparable or even superior to group-wise differences between patients and controls often reported in the literature (Da Cruz et al., 2020; Murphy et al., 2020). A dominance of microstate C, observed in the reference group and in previous resting-state reports (Bréchet et al., 2019), became clearest in the EEG-fMRI data after full correction. Overall, considering topography and temporal properties altogether, microstate B tended to show the largest differences in EEG-fMRI compared to the reference group, whereas A was the most consistent.

**Fig. 7.**
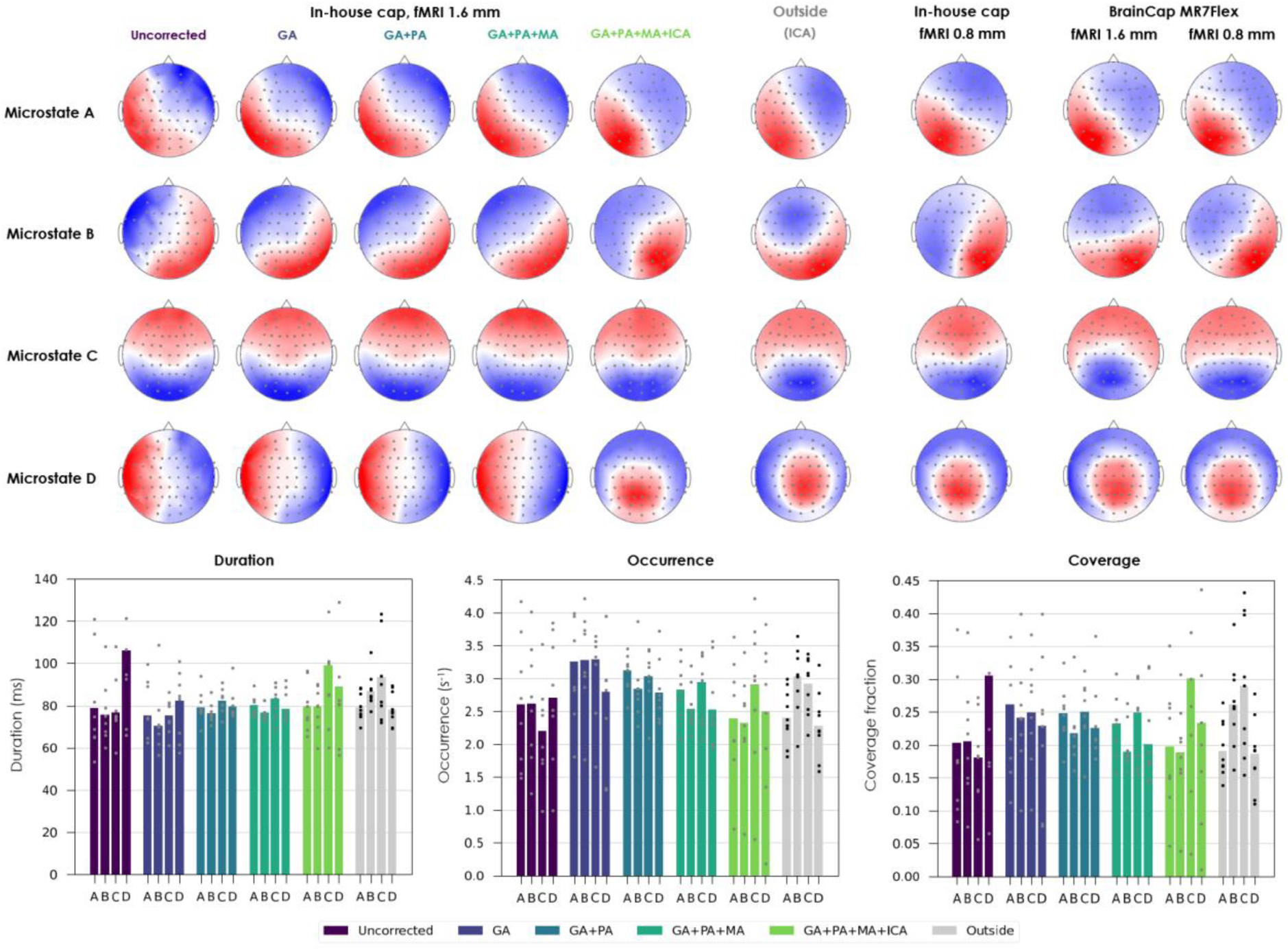
Impact of MRI-related artifacts and the respective correction steps on EEG data quality, in terms of microstate topographies and their temporal dynamics. Top left: topographies of the four microstates derived from group-level k-means clustering of the 1.6 mm EEG-fMRI data acquired with the in-house cap, estimated after each correction step. Top middle: topographies from the reference participant group recorded outside the MRI scanner, after ICA denoising. Top right: topographies obtained with the other conditions (0.8 mm fMRI, BrainCap MR7Flex), after full correction. For consistency, all cases were first downsampled to 200 Hz, bandpass-filtered to 1–40 Hz, bad channel-interpolated and re-referenced to the channel average; after clustering, for each case, the four topographies were labelled A–D based on their degree of similarity to the four canonical states typically found in resting-state EEG (Michel and Koenig, 2018). Bottom: Temporal properties of the microstates of the 1.6 mm EEG-fMRI run (in-house cap); each bar represents the average across subjects; the dot markers represent the individual subjects. The reference group is also shown, in gray, for comparison.

### 3.5. Eyes open/closing responses

The eyes-opening/closing paradigm was found to elicit robust activity changes in both fMRI and EEG data, for all the participants that underwent this task (Fig. 8, Supp. Fig. 7). On the fMRI side, ICA enabled retrieving components located on the visual cortex and oscillating in synchrony with the paradigm blocks, as expected for this type of paradigm (Jorge et al., 2015b). In some cases, the decomposition yielded more than one visual IC related to the paradigm, likely resulting in more diluted response maps (e.g. subject 2 in Fig. 8). In most subjects, additional components were also observed in higher regions of the cortex (e.g. matching the DMN configuration) that showed some degree of temporal covariation with the paradigm, suggesting an impact of the task on the activity of higher networks (Fox et al., 2005). On the EEG, every participant yielded several ICs in which the alpha power increased systematically during eyes closed, and concentrated in the occipital area (in some cases split into left and right contributions), as expected (Adrian and Matthews, 1934). The effect was generally strong enough to be directly visible on the EEG traces already after GA+PA correction, and became increasingly cleaner with additional correction steps (Supp. Fig. 8). The GLM analysis showed that the EEG-derived alpha power timecourses had a robust negative association with BOLD fMRI fluctuations in the occipital cortex for every subject (Fig. 8, Supp. Fig. 7), with peak Z-scores of −9.2 to −18.0 across subjects in the visual network (Yeo). As in the ICA maps, other activation clusters could be found in upper cortical regions as well (e.g. regions matching the DMN), yet with weaker strength and higher variability across subjects (including positive clusters in some cases). The R2adj analysis yielded peak values of fMRI variance explained by the alpha regressor of 19 to 40% across subjects. The average values over entire Yeo networks were smaller (Fig. 8); nonetheless, the alpha regressor was confirmed to be statistically meaningful, explaining significantly more variance in the visual cortex than the randomized version (*p* < 0.01).

**Fig. 8.**
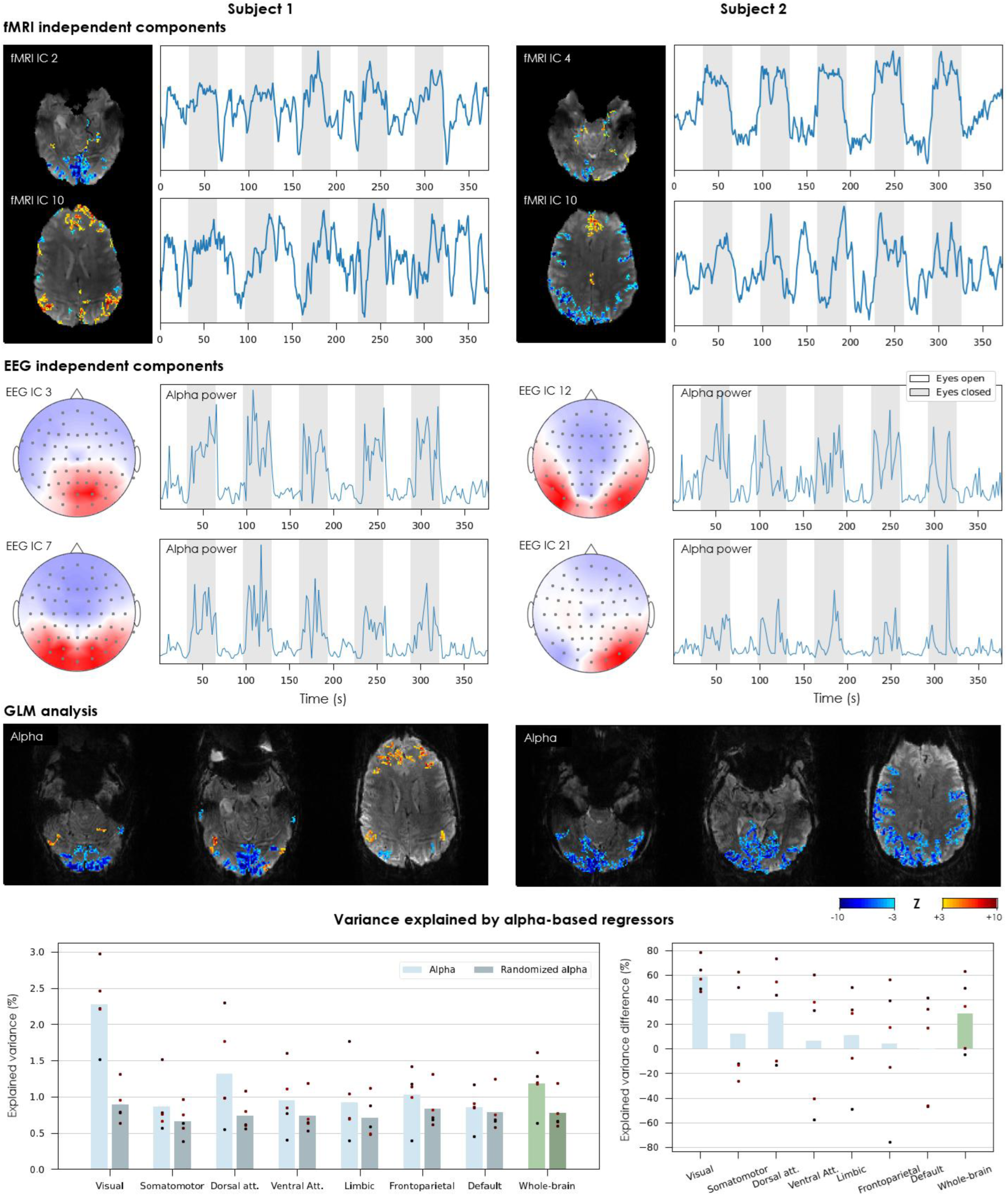
Sensitivity of simultaneously-acquired EEG and fMRI signals to activity modulation from eyes closing and opening (BrainCap MR7Flex sub-group). In the two example subjects shown, the fMRI ICs are presented in terms of their spatial distribution (converted to a Z-score map and thresholded at |*Z*| > 1.4) and timecourse. For each subject, the IC showing the strongest correlation to the paradigm is shown first, followed by a second IC associated to higher network regions that also showed some degree of co-variation with the paradigm. To harmonize with the alpha power timecourses (EEG ICs), each map-timecourse pair of fMRI ICs is displayed with a polarity such that the timecourse correlates positively with the eyes-closed blocks (i.e. high in eyes-closed, low in eyes-open). The EEG ICs are shown in terms of their scalp distribution and the timecourse of their alpha power (7–13 Hz). The two ICs showing the strongest alpha modulations locked with the paradigm are shown. The fMRI and EEG IC timecourses and the EEG IC scalp maps are shown in arbitrary units. The GLM analysis results are shown in terms of Z-score maps for the HRF-convolved EEG-derived alpha power timecourse, thresholded at |*Z*| > 3.0, in representative slices covering the visual cortex and superior cortical areas that showed significant responses. Bottom: Variance explained in different brain networks by the HRF-convolved alpha power regressor, as obtained from the EEG or after spectral phase randomization (average of 100 randomization trials). Right: relative difference in explained variance between the true and randomized alpha. Each bar represents the average across all subjects that underwent this experiment; the dot markers represent individual subjects.

## 4. Discussion

In this work, we investigated a new framework for simultaneous EEG-fMRI at 7T combining and improving upon landmark ideas from previous methodological studies, sought to enable unprecedented combined gains in fMRI and EEG signal quality, and which can be made robustly accessible to the broader neuroscience and clinical community. The framework was extensively evaluated in humans, covering safety, usability, data quality and functional sensitivity (resting-state activity and eyes-open/closed responses) on both modalities. Benefitting from the high sensitivity and acceleration capabilities of this setup, we successfully implemented and evaluated a large-FOV, sub-millimeter resolution fMRI protocol, highly relevant to emerging applications on laminar connectivity and epileptic activity propagation, for example.

### 4.1. Safety aspects

As observed for older 7T setups (Jorge et al., 2016, 2015b), the data obtained in this work suggest that GE-EPI protocols, even with SMS acceleration, are well tolerated by the current EEG-fMRI setup. Although the two tested fMRI protocols were quite different in terms of spatial and temporal sampling, and had relatively high SMS acceleration (3–4×), both remained well below the SAR safety limits of the MRI system, even running in its most conservative safety mode (Table I). The B_1_^+^_RMS_ of the full protocol remained below the recommendations of the EEG manufacturer for 3T (standard cable lengths) of 1 µT (Warbrick, 2020) – although no guidelines are currently available for 7T. The temperature measurements supported the same conclusion, with no significant heating detected during GE-EPI in any of the EEG electrodes or leads tested, in phantom or humans, with either cap (Table II).

While not affecting the electrodes or leads, the GE-EPI protocols did cause measurable heating on the EEG amplifiers when using the BrainCap MR7Flex with shorter extension cables – similar to effects we have previously observed in older 7T setups with compact EEG (Jorge et al., 2015b), and most likely caused by the gradient fields, not the RF pulses. The temperatures did not seem to reach problematic levels within the length of our sessions, but could become prohibitive in longer scans (e.g. for sleep studies). Notably, the substantially lower heating rates systematically observed for the in-house cap, and for the MR7Flex with longer extension cables, strongly suggest that opting for a slightly longer distance between the head and amplifiers may be an effective solution to minimize these heating effects (at the cost of moderate increases in EEG artifact incidence), at least for this MR gradient model.

Contrasting with these effects, in the phantom tests using SE-EPI (even within the scanner SAR limits), we did observe stronger heating along the flat ribbon strips, suggesting the formation of stronger electric currents along the leads, and thus increased safety risks. This indicates that more SAR-intensive fMRI sequences (e.g. SE-EPI, ASL, VASO) should be carefully tested, even if they obey the standard SAR restrictions. It remains unclear whether this concern also applies to the MP2RAGE sequence – the EEG recording interruptions observed with the MR7Flex with shorter cables (Table II, “Rec. stop”) happened at the scan start, pointing to its inversion RF pulse as the most likely cause, but the problem disappeared when using longer extensions. More dedicated tests should allow clarifying this, if needed for applications where the anatomical scan cannot be performed separately, without EEG. Altogether, our observations suggest that sequences employing refocusing or inversion pulses, such as SE-EPI, MP2RAGE and likely others such as ASL and VASO, should undergo more dedicated studies at 7T to better understand which regimes of operation may be safe or unsafe, and whether EEG adaptations such as longer extension cables may be consistently helpful. For users employing commercial hardware, such as commercial implementations of the MR7Flex, the respective manufacturer guidelines and sequence restrictions should naturally be the primary reference for safety aspects.

It is also important to consider the range of applicability of this study’s results. Our tests were conducted on a Siemens Terra system with a Nova 32-channel array, currently the state-of-the-art option for 7T MRI with a rapidly growing number of sites worldwide, and therefore an optimal case study in terms of applicability. However, the field continues to evolve, with parallel transmit (pTx) capabilities presenting an attractive line of exploration (as discussed below in 4.4), a new upgrade already available for the Terra (which streamlines pTx for clinical use, and optionally introduces more powerful gradients (Bell, 2024)), and other 7T models gaining traction in the market (GE Healthcare, 2021). It will be pertinent to re-visit safety aspects as new variants of the framework are explored.

### 4.2. Usability aspects

Overall, the new EEG setup did not appear to affect participant compliance (at least for this group of healthy, relatively young adults), and did not introduce major practical constraints in comparison with more established EEG-fMRI setups. The added difficulties in placement and handling of the leads inside the RF coil were substantially mitigated with practice. One difficulty that remains is in improving specific electrode contacts (e.g. adding more gel) once the subject’s head is inside the confined space of the RF coil. Therefore, it was found important to perform a good cap preparation prior to moving into the scanner, and to coach the participant to lie their head down in the coil with minimal drag. In terms of ease of access to the participant, and particularly their removal from the scanner in case of an emergency (e.g. epileptic seizure in a patient), we deem that the setup under study allows a similar approach to more conventional EEG-fMRI setups at 3T, namely by releasing the chin velcro strap of the EEG cap, detaching the ECG electrode taped to the back, and then removing the subject leaving the EEG system behind. Regarding the cleaning stage, future technical improvements to make the prototypes fully waterproof may be valuable to facilitate this task, which remained more cumbersome than with standard caps.

Future studies in larger groups, including wider age ranges and patients, will help building a more comprehensive picture of the usability of this framework, and likely reveal further opportunities for improvement. They will also help confirming the robustness of the new cap models against damage and wearing out (e.g. of the new lead segments and their interfaces); in the present study, no issues were found throughout the timespan of the experiments, on either cap prototype.

### 4.3. EEG and MRI data quality – are we there, yet?

The quality of (f)MRI data acquired with this novel framework was a central focus of investigation. The EEG cap modification recently proposed by (Meyer et al., 2020) was a landmark achievement that allowed, for the first time, to acquire EEG-fMRI at 7T with state-of-the-art imaging sensitivity and acceleration capabilities. While (Meyer et al., 2020) convincingly showed the detection of meaningful EEG and fMRI responses induced by certain cognitive tasks, the study did not characterize the level of data quality achieved by their framework, the impact of each modality on the other, and the underlying interference mechanisms. We addressed these questions with a systematic comparison of simultaneous EEG-fMRI data to unimodal data acquired in similar conditions, and extracted more transversal measures of data quality and sensitivity, as well as underlying field perturbations.

As in previous 7T setups (Jorge et al., 2015b; Lê et al., 2022), our tests indicated that the disruption of the B_1_ field (i.e. RF pulses) is the main cause of MRI data degradation, whereas B_0_ is not substantially affected inside the brain (Fig. 2, Supp. Fig. 1, Fig. 3). Importantly, the severity of the B_1_ perturbations was found substantially milder than in older 7T setups, and without focal signal losses, which appears in agreement with the qualitative observations of (Meyer et al., 2020) for their setup. This improvement may originate from the change in RF coil and/or from the EEG cap modification; one strategy to distinguish between the two effects would be to run the same framework while changing only the coil, but unfortunately our Terra platform did not have an alternative head coil available to test. Nonetheless, while the source of the improvement could not be fully understood, its benefits were consistent across the participant group, with remarkably moderate SNR losses across the brain (Fig. 3, Fig. 4). Another favorable observation was that the most affected regions, which tended to be the occipital, parietal and frontal lobes, were also those benefitting from the highest SNR without EEG – leading to a more balanced distribution with EEG. Importantly, the 0.8 mm protocol had higher losses than the 1.6 mm – this may be linked to its higher level of undersampling, which might make it more sensitive to the accuracy of the parallel imaging calibration, and therein more extensively affected by B_1_ penalties. Of note, an additional analysis carried out on the 0.8 mm data to investigate the influence of partial volume effects on the parcellations for the different resolutions under study (namely 1.6 mm vs. 0.8 mm GE-EPI) indicated that partial volume effects did not contribute significantly to the differences in MRI quality and functional sensitivity observed between these two fMRI protocols (Supp. Fig. 9).

Interestingly, while both EEG cap prototypes induced relatively similar effects on the MRI data, the MR7Flex did consistently show a milder impact than the in-house prototype on essentially every metric, and particularly on the temporal SNR of both fMRI protocols, which is a critical measure for functional imaging. While the data does not allow confirming the reasons for this outcome, a critical difference between the two caps is that the MR7Flex contains point resistors segmenting the leads relatively close to the electrodes (on the four PCBs – Fig. 1-10). We have previously found that segmenting the EEG leads in this manner may substantially mitigate their interactions with RF pulses (Lê et al., 2022). Another potential contribution relates to the amount of conductive cabling and its distribution: in the MR7Flex, the tinsel leads do not all extend all the way to the same point on the scalp or form a large lead bundle; instead, they converge to four separate PCBs, and then follow on the thinner tracks of the ribbon strips (Fig. 1). In any case, the superior performance of the BrainCap MR7Flex is a positive finding, as this option has the highest readiness level towards more widespread distribution.

Another encouraging observation was the decreasing impact of EEG when moving from spatial to temporal SNR, and finally to the diverse functional sensitivity metrics tested here. In both fMRI protocols, the reductions in temporal SNR due to EEG tended to be milder than those in spatial SNR (Fig. 4). This non-linear relationship between spatial and temporal SNR has been observed in diverse other contexts (Triantafyllou et al., 2005), and may be influenced by contributions such as so-called physiological noise (Van Der Zwaag et al., 2015), which scales with signal strength (Krüger and Glover, 2001), and may thus impose a smaller burden on the temporal SNR when the spatial SNR is reduced by EEG (Luo and Glover, 2012). Further along, considering the fALFF, INC and pair-wise functional connectivity, their natural variability across subjects and sessions seems to have superseded any eventual systematic effects from the introduction of EEG – despite behaving quite consistently in other aspects that indicated functional sensitivity (Fig. 5, Supp. Fig. 2 and Supp. Fig. 3, Results section 3.3). Likewise, although certain brain regions seemed more affected by EEG than others in terms of spatial SNR, this effect did not stand out in the functional analyses, even in more detailed cortical parcellations such as the DK atlas (Supp. Fig. 4). The ICA results also agreed with these observations, with the resting-state fMRI data (both protocols) yielding similar networks with and without EEG (Fig. 5), and the eyes-open/closing data yielding occipital network responses locked to the paradigm in every tested participant (Fig. 8). Likewise, the GLM analysis revealed that EEG alpha power was robustly associated to fMRI fluctuations in the visual cortex, in line with previous studies at 3T (Goldman et al., 2002; Ingram et al., 2024), for every tested subject.

In general, although the spatial and temporal SNR losses observed in this work did not translate into systematically measurable losses in resting-state functional sensitivity or connectivity, and did not preclude the detection of strong alpha-related BOLD fluctuations, those penalties in SNR may nonetheless exert more important effects in other applications. Of note, the SNR losses reported in this work for different brain regions and networks may offer practical insights to help planning future EEG-fMRI studies, within their range of applicability. For example, if a user obtains pilot fMRI data for their application of interest, and their fMRI protocol does not widely differ from the ones tested here, then they can use our SNR values and % losses as a reference, to understand the expected impact that adding EEG may have on the contrast-to-noise ratio (CNR) of the functional responses they are targeting, or on the distinguishability of their functional differences – and hence better understand whether the resulting compromises in detection power may be acceptable for their use-case or not.

Regarding EEG quality, an important observation was that saturation effects were relatively rare, and thus did not pose a bottleneck to EEG recording in this setup, even during 0.8 mm-resolution fMRI– indicating excellent prospects for high-resolution imaging, one of the strongest advantages offered by 7T. Still, the saturation effects that did occur in our dataset were caused by GA peaks and appeared more predominantly at 0.8 mm, suggesting that this possibility should be given particular attention when preparing for very high-resolution applications. As no saturation preference was found for specific EEG channels, with either cap prototype, it is unclear whether modifications to the lead arrangements could be helpful; once again, promoting good cap preparations with low contact impedance may be the best strategy to invest in altogether.

The different EEG artifact contributions observed in this work were in line with previous reports (Debener et al., 2008; Jorge et al., 2015a; Neuner et al., 2014). These artifacts could supersede the activity of interest by several orders of magnitude, obscuring even the strongest marks of healthy activity such as the alpha peak, and spanning a wide range of frequencies of interest (Fig. 6, Supp. Fig. 5). However, through the use of effective correction approaches (Warbrick, 2022), including the use of reference sensors (Jorge et al., 2015a; Warbrick and Schubert, 2020), it was found possible to reduce the artifacts to a large extent. The resulting signals of this participant group were then mostly equivalent to the reference group recorded outside the scanner, in terms of spectral profile, dominant topographies, and their temporal properties (Fig. 6, Fig. 7). Of note, the microstate analysis indicated more pronounced deviations with respect to the reference group for certain microstates (namely B; Fig. 7) than others (namely A). Such differences could be due to residual EEG artifacts, and/or to the removal of true neuronal activity by the artifact correction methods, which always carry certain trade-offs between correction efficacy and neuronal signal preservation. At the same time, natural variability in large-scale neuronal activity may have played a role in the observed differences as well. Previous work looking at test-retest variability of microstates within and across sessions found that reliability is higher toward the start of the sessions, and appears to covary to some extent with subjective ratings of alertness; at rest (mind wandering), microstate B in specific was found to have a relatively poor reliability in terms of coverage and occurrence estimates (Antonova et al., 2022). Since our EEG-fMRI and reference groups underwent different experimental procedures (in particular, the EEG-fMRI group underwent an MRI-only session beforehand), it is plausible that there may have been systematic differences in alertness and fatigue, which could contribute to the observed differences as well, namely with microstate B. In parallel, the evidence obtained from the eyes-open/closing runs (Fig. 8, Supp. Fig. 7, Supp. Fig. 8) indicates excellent functional sensitivity for alpha modulation. Future studies and more diverse applications will be helpful to expand and clarify the current limits of applicability.

### 4.4. Opportunities for further improvement and validation

On the MRI side, an interesting finding from the B_1_ data is that, even in the conventional case without EEG, the transmit voltage indicated by the scanner’s standard calibration systematically led to an average B_1_^+^ across the brain below 0.8 (according to our SA2RAGE maps; Fig. 3), where 1.0 was the target outcome. While this did not invalidate any comparisons between without and with-EEG conditions in the dataset, it suggests that the transmit calibration approach may benefit from dedicated improvements, to better approximate the desired (nominal) flip angles and maximize SNR. Our current observations on SAR and heating do suggest this could be accommodated by the setup. Also regarding B_1_, it was found that the adjustment (increase) in transmit voltage proposed for the with-EEG cases did consistently improve the transmit field (Fig. 3), although only partially recovering the EEG-induced B_1_^+^ losses, and without curbing the increase in spatial heterogeneity. This is a reasonable outcome since we employed single-channel transmission, and strongly motivates the exploration of more flexible pTX techniques, to recover both B_1_ strength and homogeneity – especially as the perturbations from this new EEG setup are relatively smooth, not highly localized. On the other hand, the improvements observed in GRE image SNR when increasing the transmit voltage were more moderate than for transmit B_1_ (Fig. ^3^). This may be a consequence of the fact that the EEG affects both the transmit and receive fields; both contribute to SNR, and the receive side will still be affected even if the desired flip angle is fully achieved. Thus, to close this last gap in MRI data quality, additional improvements may still prove valuable, such as with new, higher-resistivity lead materials (Levitt et al., 2023; Poulsen et al., 2017).

In terms of fMRI specificity, while the present work was more oriented towards pushing spatial resolution, it could likewise be interesting to explore the temporal dimension in future work, with very fast fMRI protocols. As far as we know, all EEG-fMRI studies that have explored SMS-EPI so far, at 7T (Jorge et al., 2019, 2016) and 3T (Chen et al., 2020; Egan et al., 2021; Uji et al., 2018), have tested acceleration factors up to 4×, similar to our 1.6 mm protocol, achieving shorter TRs mainly by setting thicker slices or slice gaps, to cover the same FOV with less slices. We hypothesize that a similar approach could be explored with our setup, and would likely result in even milder heating effects over time, given the lower gradient demands in this case. On the other hand, the high image quality observed with the 1.6 mm protocol at 4× SMS acceleration suggests that it could be interesting to further push the acceleration factor, without substantially dropping the spatial resolution. In this case, the protocol would likely benefit from more dedicated testing to evaluate its heating effects over time.

Regarding EEG quality, the artifact correction procedures applied in this work could potentially benefit from more dedicated methodological testing, particularly to reduce their manual input needs. The current pipeline was built upon our previous results at 7T (Jorge et al., 2019, 2015a; Wirsich et al., 2021), and focused on achieving the best possible data quality, at the cost of extensive manual inputs in (i) reviewing cardiac triggers, and especially (ii) reviewing the ICA decompositions to optimally identify artifactual ICs. The need for cardiac triggers could potentially be obviated by relying fully on the artifact sensor signals to address both PA and MA contributions together. The ICA step was found to contribute with relevant improvements to the data (Fig. 6, Fig. 7, Supp. Fig. 8), suggesting it should not be skipped, at least for some applications; alternatively, it may be possible to expedite the process with automatic selection criteria (Abreu et al., 2016; Mantini et al., 2007). Another possibility that would be interesting to explore would be replacing ICA with a beamforming approach, which performs EEG source localization by means of a spatial filter that also reduces artifact contributions (Brookes et al., 2008; Uji et al., 2021); this approach requires less manual input than ICA, although the electrode positions need to be accurately known. Additionally, to better clarify the potential contributions to the microstate differences observed between the EEG-fMRI and outside-MRI groups (Fig. 7), it would be of significant interest to consider a dedicated cohort study covering test-retest reliability within and across sessions, including both EEG-fMRI and EEG-only sessions.

As this acquisition framework becomes more widely available to the research community, further assessments of quality and functional sensitivity for other functional features will be pertinent as well. The EEG quality metrics explored here cover a broad range of information contained in EEG data: oscillations occurring in various bands within 0.75–70 Hz have been a major source of meaningful cognitive information in neuroscience studies, both basic and clinical (Niedermeyer and Lopes da Silva, 2005), and microstate representations have been found sensitive to alterations across diverse cognitive states and disorders (Tarailis et al., 2024). Nonetheless, other major EEG features such as event-related potentials will benefit from more focused tests, for instance to evaluate sensitivity and accuracy when estimating parameters such as latency, component amplitudes, and their variability across trials (Jorge et al., 2019). On the clinical side, major applications such as epilepsy will benefit from more targeted assessments of interictal spike detection, for instance (van Graan et al., 2015). Regarding fMRI sensitivity, it is important to note that the tests and analyses performed in this study were mainly centered on resting-state activity, and on responses associated with alpha power modulation (eyes open/closed). While the results indicate very positive prospects in terms of fMRI sensitivity over the whole brain, it will be important to pursue dedicated task-based studies to more thoroughly assess the impact of EEG on fMRI responses in specific regions, such as the visual and sensorimotor cortices, which are of wide interest to neuroscience research. This may be particularly pertinent to confirm the feasibility of specific layer-based analyses, for example, given their challenges in terms of SNR.

## 5. Conclusions

The novel EEG-fMRI framework investigated in this work shows an overwhelmingly positive performance in terms of safety, usability and data quality. With this setup and recording parameters, simultaneous acquisitions, even with sub-millimeter fMRI resolution, could be conducted without detectable safety issues or major practical constraints. Relatively mild penalties were found in MRI quality, without detectable impairments in functional sensitivity to resting-state fluctuations and eyes-open/closed responses. EEG artifacts could be extensively mitigated by optimized correction techniques, rendering the signal characteristics largely comparable to recordings outside MRI. While further opportunities for technical improvement have been identified, and more dedicated validations in task-based conditions need to be pursued, the current evidence indicates excellent prospects for neuroimaging applications using this framework, that can leverage the unique possibilities offered by 7T functional imaging.

## Acknowledgments & funding information

This work was funded by the Swiss National Science Foundation through grant PZ00P2_185909, and supported by CSEM – Swiss Center for Electronics and Microtechnology, by the Translational Imaging Center (TIC) of the Swiss Institute for Translational and Entrepreneurial Medicine (SITEM), and by CIBM Center for Biomedical Imaging, Switzerland. We thank Roberto Rusconi for his assistance in the modifications of the in-house EEG cap.

## Data availability

The human EEG-fMRI dataset collected in this work, including both fMRI protocols and both EEG caps, has been made publicly available in Zenodo, at https://doi.org/10.5281/zenodo.15781280. The data are provided after artifact correction and pre-processing, together with anatomical and functional brain parcellations, and are ready to undergo resting-state analyses.

## Author contributions (CRediT)

**CS Martinez:** Conceptualization, Methodology, Software, Validation, Formal analysis, Validation, Investigation, Data curation, Writing – review & editing, Visualization, Project administration. **J Wirsich:** Conceptualization, Methodology, Investigation, Resources, Writing – review & editing. **C Jäger**, **T Warbrick, J Bastiaansen:** Conceptualization, Methodology, Resources, Writing – review & editing. **S Vulliémoz**, **M Lemay**, **R Wiest:** Conceptualization, Resources, Writing – review & editing. **J Jorge:** Conceptualization, Methodology, Software, Validation, Formal analysis, Investigation, Resources, Data curation, Writing – original draft, Writing – review & editing, Visualization, Supervision, Project administration, Funding acquisition.

## Supplementary Figures

**Supp. Fig. 1.**
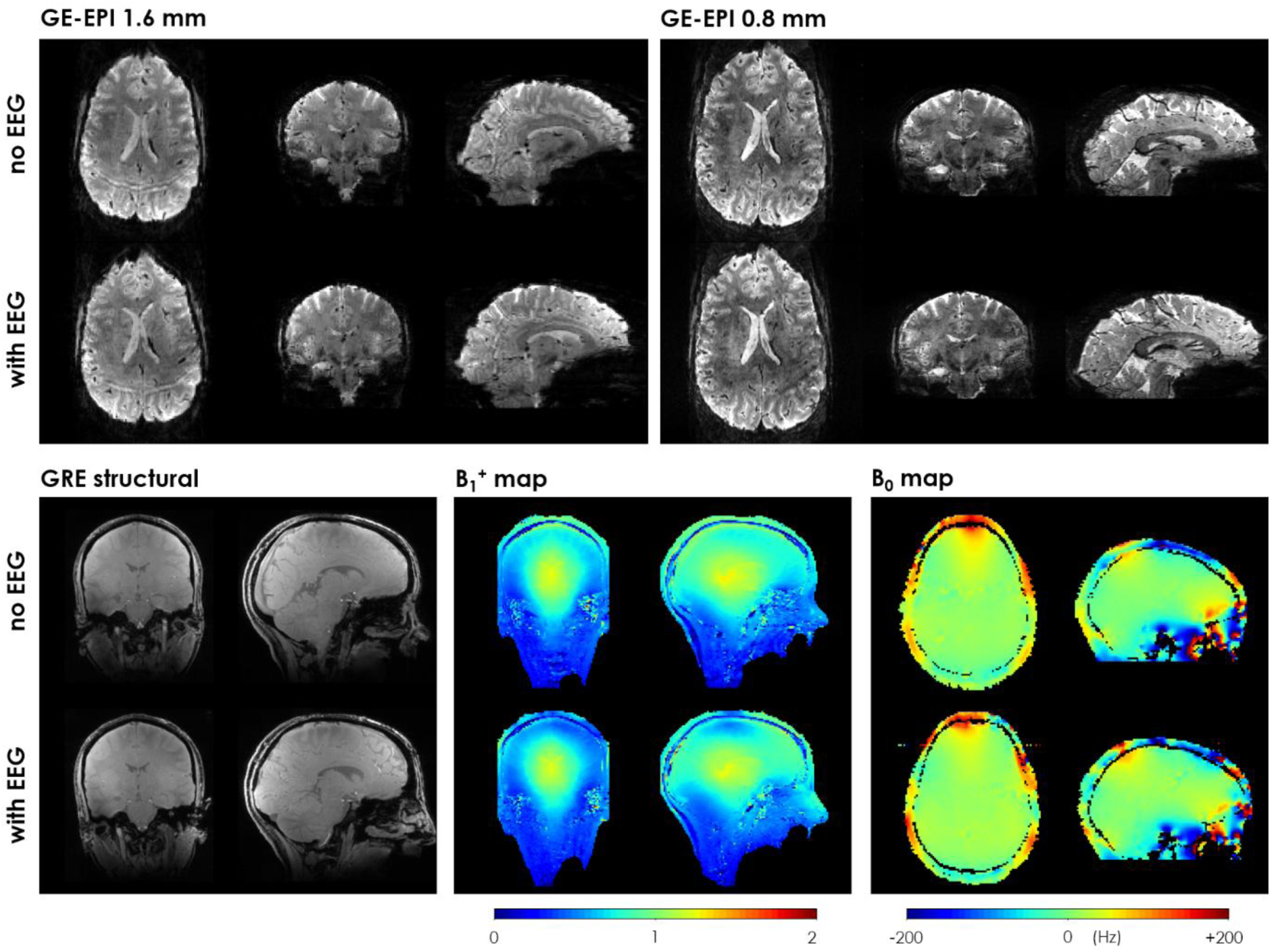
MRI data obtained from an example subject with and without the EEG system in place. This example used the BrainCap MR7Flex prototype. The slices shown include some of the most relevant differences found between the with- and without-EEG conditions. Top: GE-EPI volumes from the fMRI acquisitions (left: 1.6mm isotropic resolution; right: 0.8 mm isotropic resolution). Bottom: GRE-based anatomical image and field maps. The B_1_^+^ map is expressed as a fraction of the nominal flip angle. Both maps were masked to exclude background voxels.

**Supp. Fig. 2.**
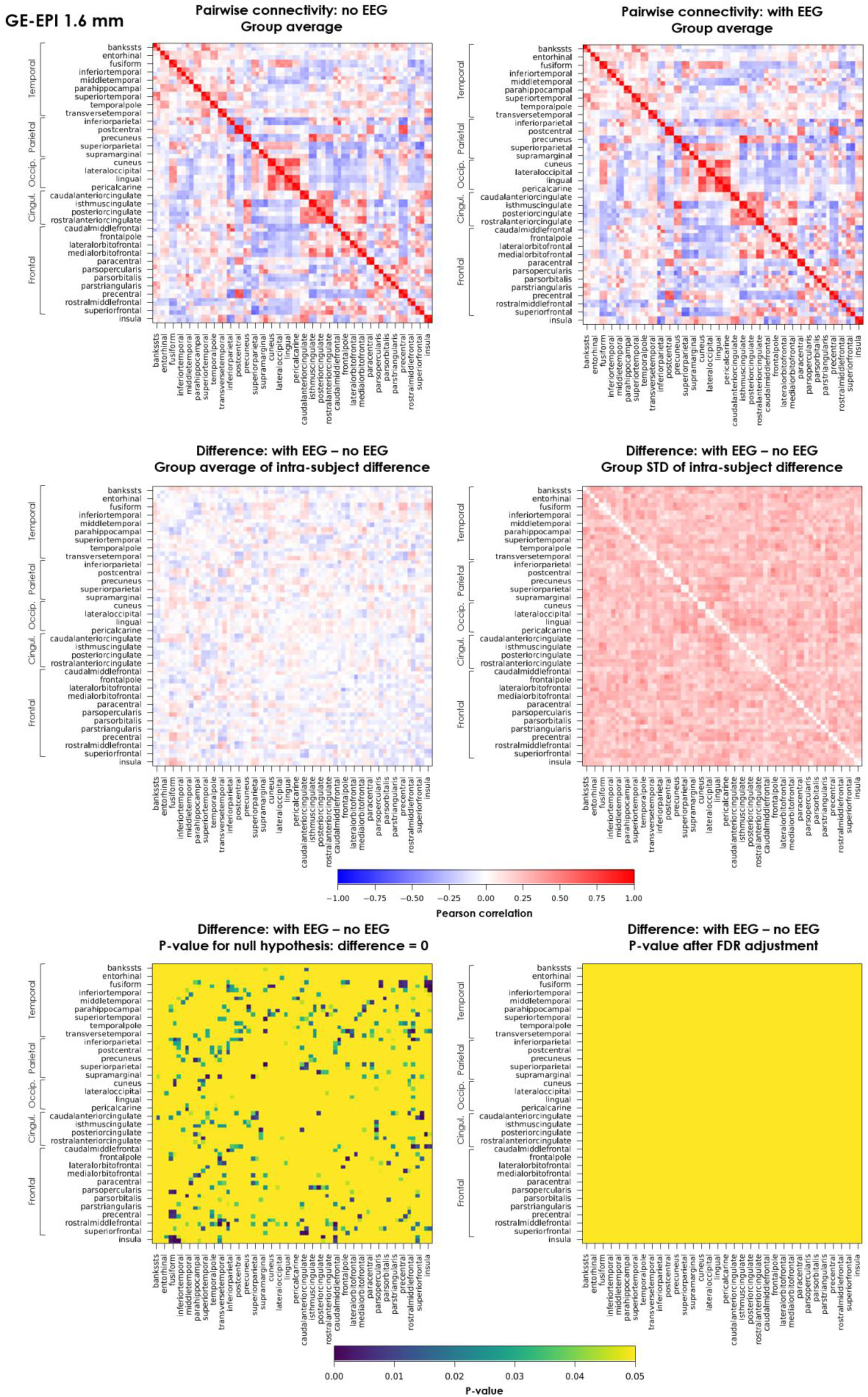
Impact of EEG on fMRI resting-state connectivity across different cortical region pairs (DK atlas), evaluated on the 1.6 mm GE-EPI acquisitions with the in-house EEG cap prototype. Top row: functional connectivity matrix, averaged across subjects, for the acquisitions without and with EEG. Prior to group averaging, the individual matrices were obtained following the approach described in (Wirsich et al., 2021), based on Peason correlation. Each region label is centered on a pair of rows (resp. columns) which refer to the left and the right hemisphere for the same region. Middle row: group average (left), and group standard deviation (right) of the difference matrices between without- and with-EEG acquisitions of each subject. Bottom row: p-values for the group-level statistical significance of a difference in functional connectivity between without and with EEG conditions, for each DK region pair; the p-values are shown without correction for multiple comparisons on the left, and after adjustment based on false discovery rate (FDR) (Benjamini and Hochberg, 1995) on the right (*false_discovery_control* function from SciPy, Python). The color scale is given an upper limit of *p* = 0.05, as a threshold for significance.

**Supp. Fig. 3.**
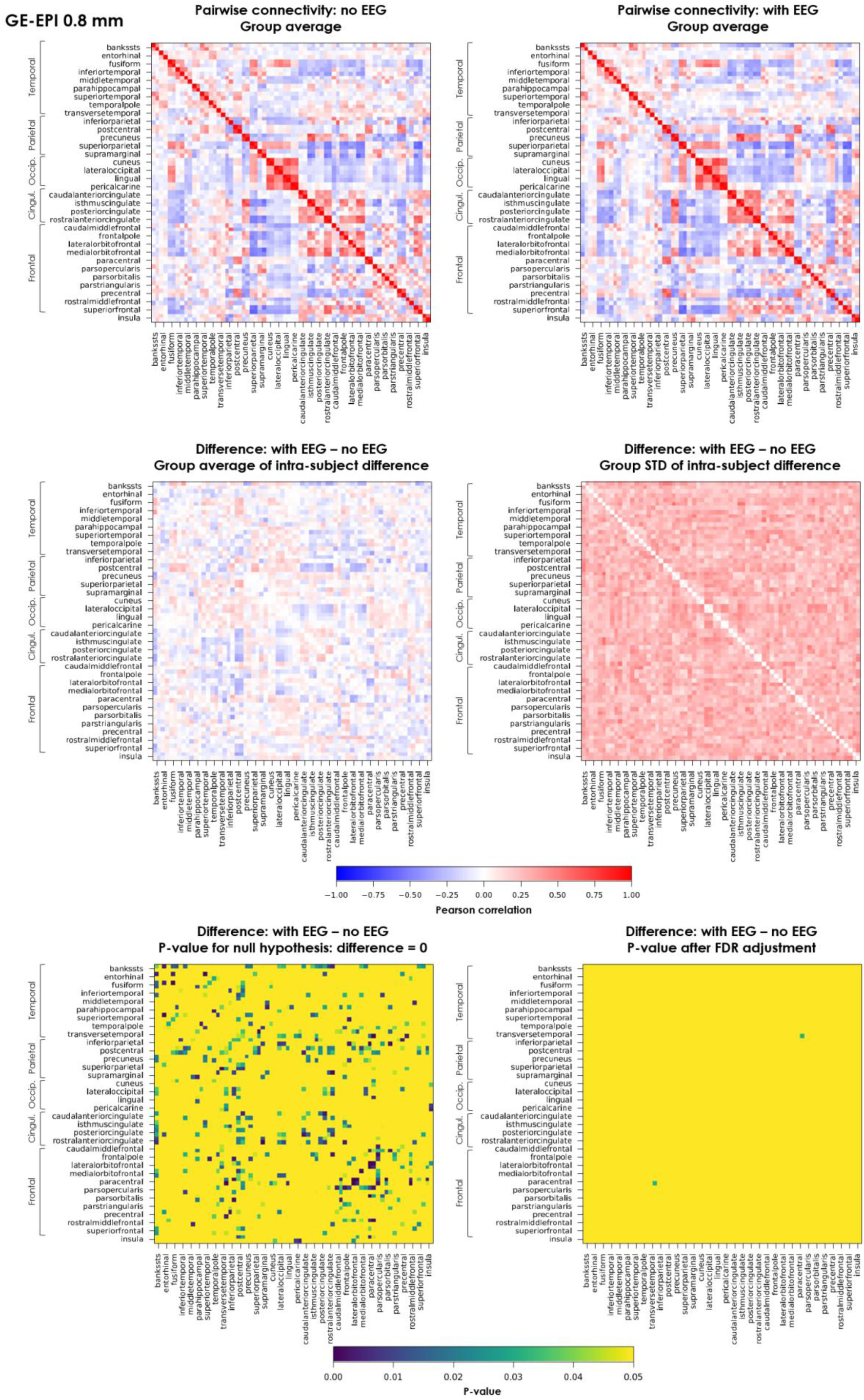
Impact of EEG on fMRI resting-state connectivity across different cortical region pairs (DK atlas), evaluated on the 0.8 mm GE-EPI acquisitions with the in-house EEG cap prototype. Top row: functional connectivity matrix, averaged across subjects, for the acquisitions without and with EEG. Prior to group averaging, the individual matrices were obtained following the approach described in (Wirsich et al., 2021), based on Peason correlation. Each region label is centered on a pair of rows (resp. columns) which refer to the left and the right hemisphere for the same region. Middle row: group average (left), and group standard deviation (right) of the difference matrices between without- and with-EEG acquisitions of each subject. Bottom row: p-values for the group-level statistical significance of a difference in functional connectivity between without and with EEG conditions, for each DK region pair; the p-values are shown without correction for multiple comparisons on the left, and after adjustment based on false discovery rate (FDR) (Benjamini and Hochberg, 1995) on the right (*false_discovery_control* function from SciPy, Python). The color scale is given an upper limit of *p* = 0.05, as a threshold for significance.

**Supp. Fig. 4.**
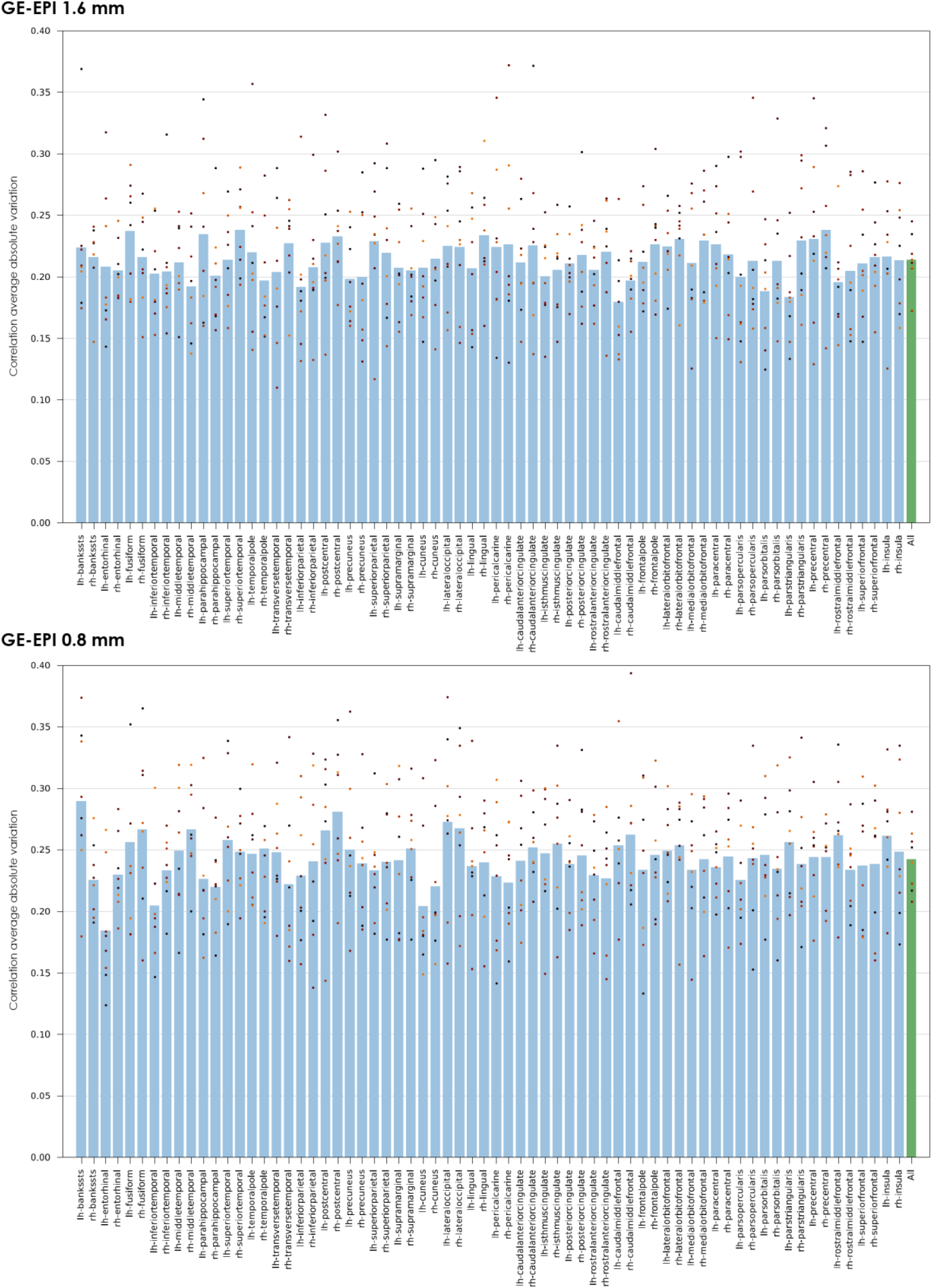
Impact of EEG on fMRI resting-state connectivity across different cortical regions (DK atlas), evaluated on the 1.6 mm and 0.8 mm GE-EPI acquisitions with the in-house EEG cap prototype. Each dot marker for each DK region represents the absolute difference in Pearson correlation between without- and with-EEG conditions, averaged across all pairs that include that region, for one individual subject; the bar represents the respective average across subjects. The Pearson correlation estimates were obtained following the approach described in (Wirsich et al., 2021). In the region labels, “lh” and “rh” indicate the left and right hemisphere, respectively.

**Supp. Fig. 5.**
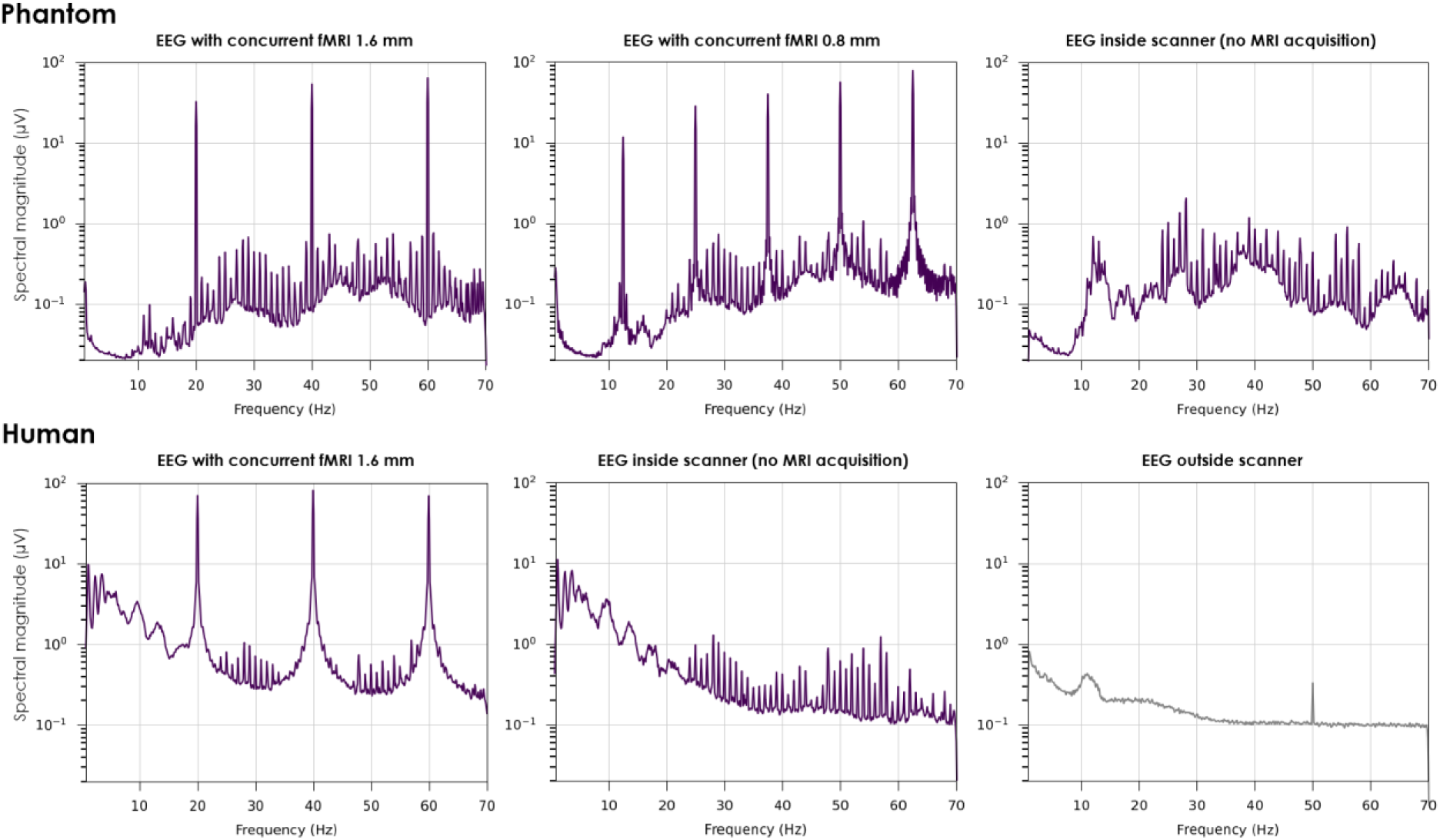
Impact of different MRI-related artifacts on EEG recordings, in a phantom (top) and an example participant (bottom), with and without concurrent fMRI acquisitions, as well as outside the scanner in a non-shielded room (for the human participant). For consistency with the conditions shown in Fig. 6, the source EEG recordings were first downsampled to 200 Hz, bandpass-filtered to 0.75–70 Hz and re-referenced to the channel average. Magnitude spectra were then estimated using Welch’s method (10 s Hann window, 50% overlap) for each channel, and then averaged across channels. For reference, the slice GA peaks are expected at 20 Hz for the 1.6 mm fMRI protocol (respectively 12.5 Hz for the 0.8 mm protocol) and harmonics; the volume GA peak, if present, is expected at 0.95 Hz (respectively 0.28 Hz).

**Supp. Fig. 6.**
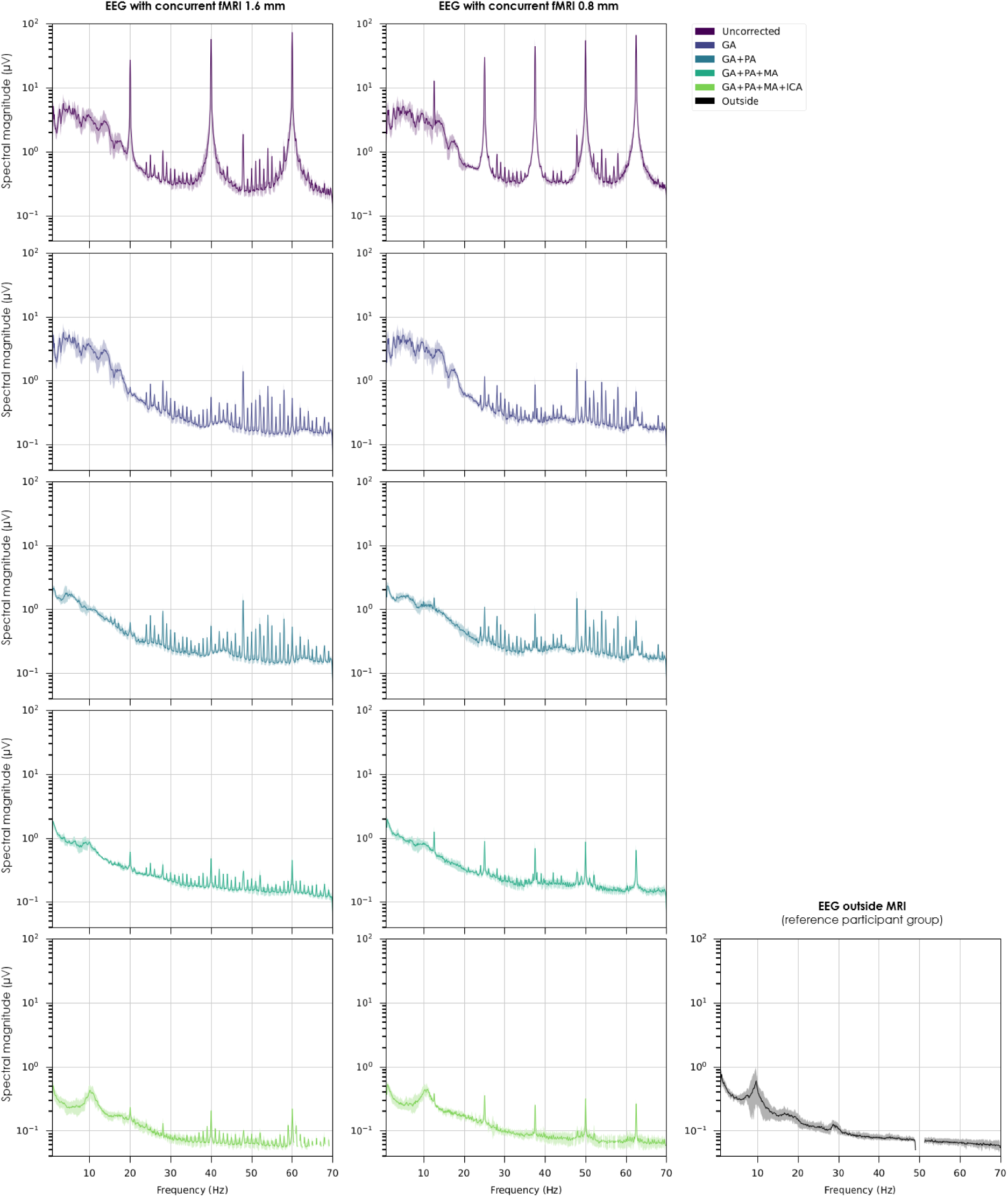
Impact of MRI-related artifacts and the respective correction steps on EEG data quality, in terms of spectral content. Left and middle column: signal spectra for the EEG-fMRI group, with either fMRI protocol, after different correction steps; for consistency, all cases were first downsampled to 200 Hz, bandpass-filtered to 0.75–70 Hz and re-referenced to the channel average; magnitude spectra were then estimated using Welch’s method (10 s Hann window, 50% overlap) for each channel, and averaged across channels. The colored lines represent the average of these signals across participants; the colored error margins represent the STD across participants. For reference, the slice GA peaks are expected at 20 Hz for the 1.6 mm fMRI protocol (respectively 12.5 Hz for the 0.8 mm protocol) and harmonics; the volume GA peak, if present, is expected at 0.95 Hz (respectively 0.28 Hz). Bottom-right: signal spectrum for the reference participant group recorded outside MRI (non-shielded room), pre-processed in a similar manner, except for an additional 50 Hz notch filter; the black line denotes the average across participants; the gray error margin denotes the STD across participants.

**Supp. Fig. 7.**
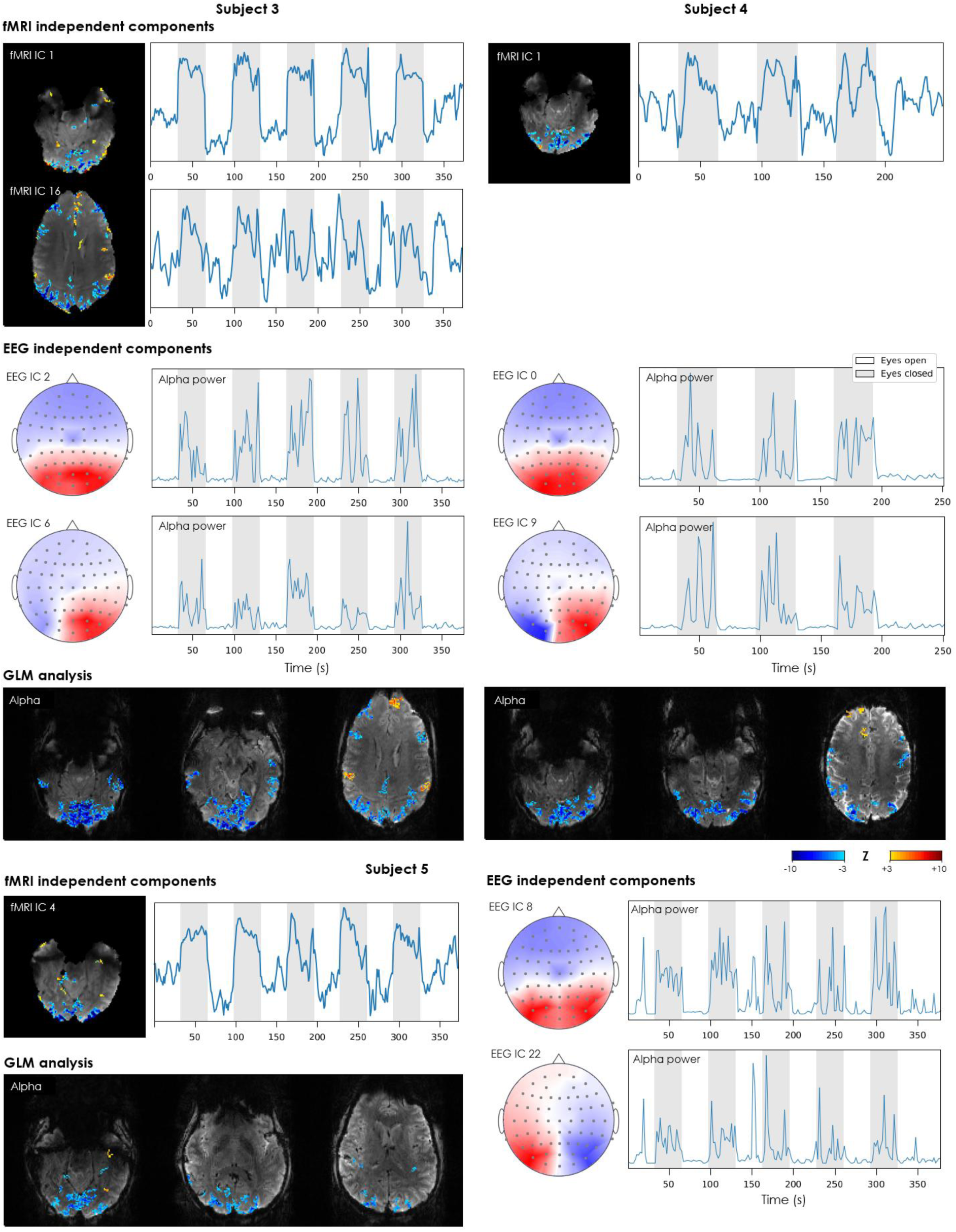
Sensitivity of simultaneously-acquired EEG and fMRI signals to activity modulation from eyes closing and opening, complementing Fig. 8 with the other 3 subjects that underwent this task (BrainCap MR7Flex sub-group). The fMRI ICs are shown in terms of their spatial distribution (converted to a Z-score map and thresholded at |*Z*| > 1.4) and timecourse. To harmonize with the alpha power timecourses (EEG ICs), each map-timecourse pair of fMRI ICs is displayed with a polarity such that the timecourse correlates positively with the eyes-closed blocks (i.e. high in eyes-closed, low in eyes-open). The EEG ICs are shown in terms of their scalp distribution and the timecourse of their alpha power (7–13 Hz). The two ICs showing the strongest alpha modulations locked with the paradigm are shown. The fMRI and EEG IC timecourses and the EEG IC scalp maps are shown in arbitrary units. The GLM analysis results are shown in terms of Z-score maps for the HRF-convolved EEG-derived alpha power timecourse, thresholded at |*Z*| > 3.0, in representative slices covering the visual cortex and superior cortical areas that showed significant responses.

**Supp. Fig. 8.**
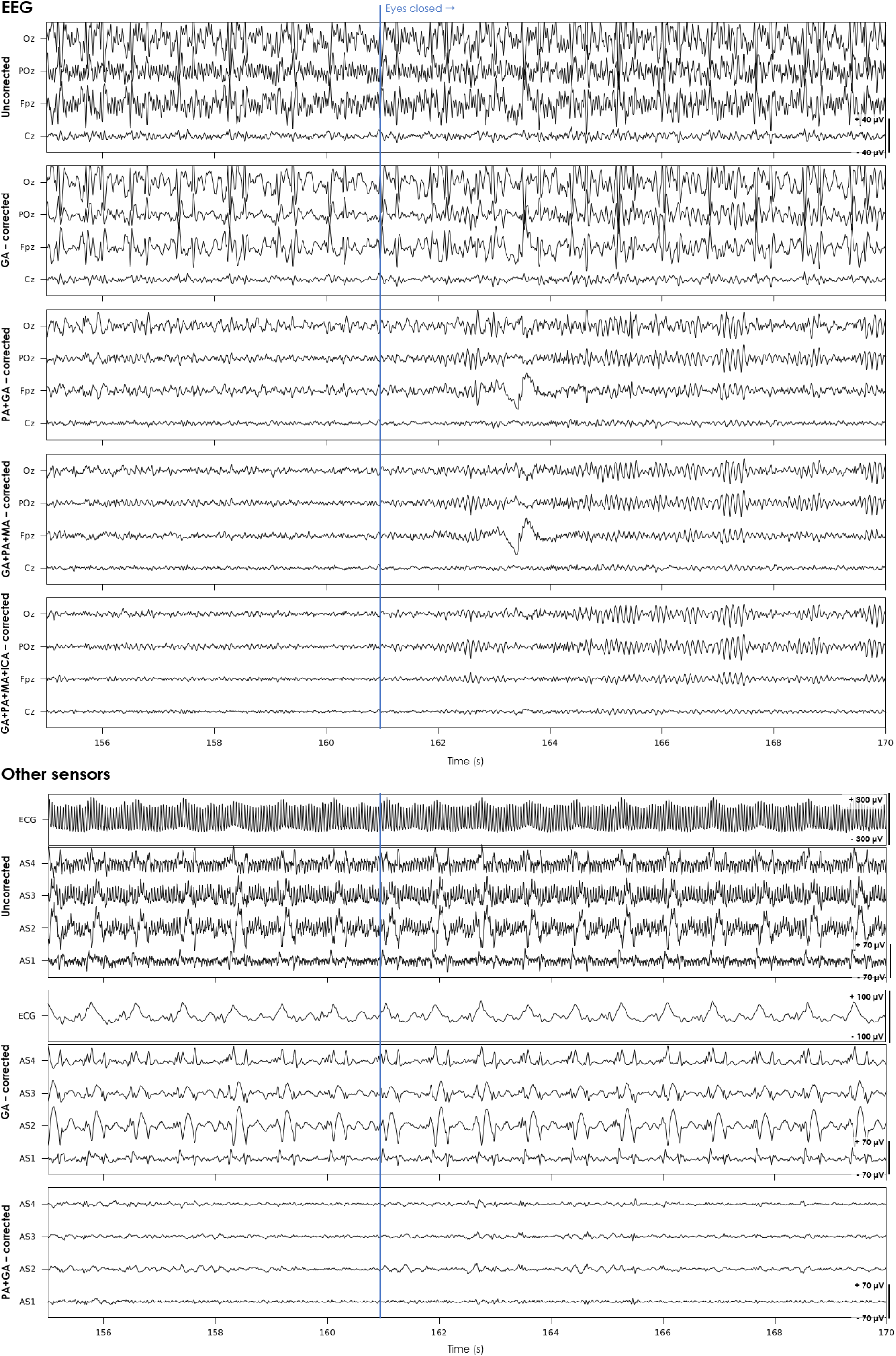
Impact of MRI-related artifacts and correction steps on EEG traces (top) and other sensors (ECG and artifact sensors; bottom), shown for an example period from an eyes-open/closing run (concurrent 1.6 mm fMRI, BrainCap MR7Flex). The vertical blue line marks a queue to close the eyes. For consistency, all cases are shown after downsampling to 200 Hz, bandpass-filtering to 1–40 Hz, bad channel interpolation and re-referencing to the channel average; all EEG traces are plotted with the same amplitude scale (−40 to +40 µV). As can be observed, the uncorrected EEG signals are dominated by high-frequency artifacts on most channels; after GA correction, PA epochs become the dominant feature; after PA correction, the increase in alpha waves with the eyes closed now becomes readily visible in occipital channels (Oz, POz); nonetheless, the MA and ICA-based correction steps still bring improvements, reducing artifacts such as probable PA residuals and the large deflection observed in Fpz near the transition moment (likely resulting from eye movements). The fully corrected run exhibits visibly clean traces with well-preserved alpha waves, which are strongly amplified when the eyes are closed, particularly in occipital channels. We note that the ICA-based correction step performed here followed the same criteria as for the resting-state runs (described in section 2.4.1, Supp. Table II), and did not focus on selecting specifically alpha-related ICs. In addition to EEG, the other sensors are displayed up until the step where they are used for EEG correction.

**Supp. Fig. 9.**
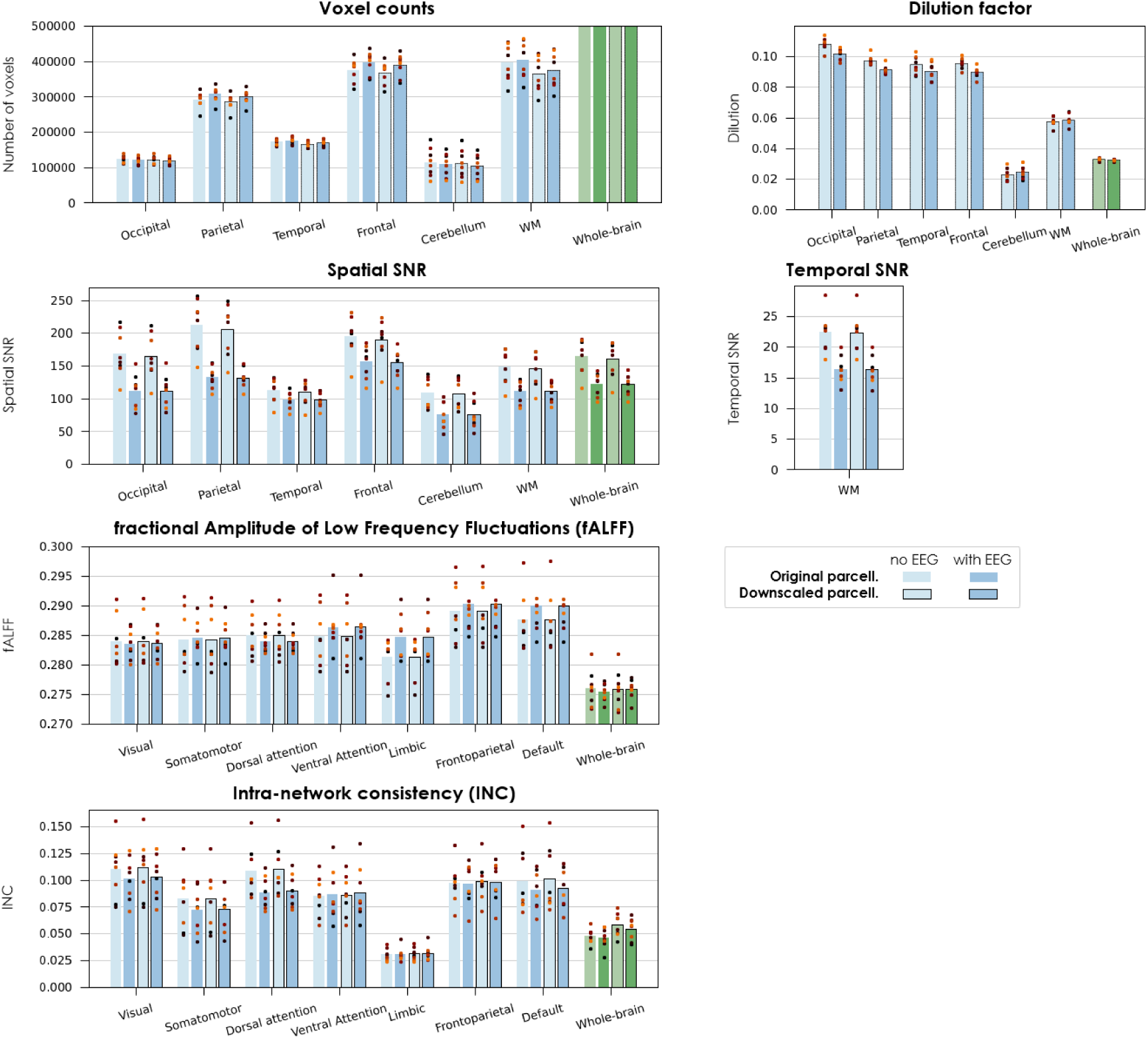
Impact of partial volume effects on the evaluation of MRI data quality without and with EEG, for different brain regions and networks. This analysis was focused on the 0.8 mm GE-EPI data, with the in-house EEG cap, and covered the main metrics for image quality and functional sensitivity considered in the study. To test the influence of partial volume effects, each of the parcellations that had been defined for the 0.8 mm data was “artificially downscaled”: the grid was split in cubes of 2×2×2 voxels, and for each cube, all voxel labels were replaced by the majority label of the cube. In this way, the parcellation became “coarser”, to a degree comparable to the 1.6 mm data and hence with similar partial volume effects, while preserving other parameters such as the voxel grid, spatial resolution and noise properties of the 0.8 mm data. The main quality metrics were then computed for these downscaled parcellations (bars with black contour), and compared with the original results (bars without contour). Two additional metrics were computed as well: (i) the number of voxels in each brain region, to assess whether certain regions tended to gain or to lose voxels in the coarser parcellation; and (ii) a “dilution factor”, computed as 1 minus the average value in each downscaled region (coarse parcellation) of an image where every voxel inside the original region (finer parcellation) is assigned the value of 1, and every voxel outside is assigned 0 – hence, the larger the amount of 0’s included, and 1’s excluded, in the coarse region, the higher the resulting dilution value. Each bar represents the average across subjects; the dot markers represent the individual subjects. Optimized EEG-fMRI at 7T for highly-resolved brain imaging

## Supplementary Tables

**Supp. Table I.** MRI sequences and parameters used in this study Optimized EEG-fMRI at 7T for highly-resolved brain imaging.

| Type | Sequence | Timings (ms) | Flip angle (°) | Resolution<sup>a</sup> (mm) | Field of view (mm) | Readout bandwidth (Hz/Px) | Undersampling | Acq. time (min) |
| --- | --- | --- | --- | --- | --- | --- | --- | --- |
| Structural<br>Low SAR | 3D GRE | TE/TR = 3.5/10 | 6 | 1.0 | 220×166×192<br>(axial slab) | 240 | 2×2 GRAPPA<br>7/8×7/8 partial Fourier | 1.3 |
| Structural<br>T<sub>1</sub>-weighted | 3D CS MP2RAGE<br>(Mussard et al., 2020) | TE/TR = 2.1/6000<br>TI<sub>1</sub>/TI<sub>2</sub> = 800/2700 | 4/5 | 0.6 | 240×226×154<br>(sagittal slab) | 240 | 4.0× on the PE<sup>b</sup> plane<br>252 samples/TR | 7.3 |
| B<sub>0</sub> mapping | Double-echo 2D GRE | TE<sub>1</sub>/TE<sub>2</sub>/TR =<br>3.0/5.2/20 | 8 | 2.0 | 220×168×144<br>(72 axial slices) | 630 | 2× GRAPPA | 0.65 |
| B<sub>1</sub> mapping | 3D SA2RAGE<br>(Eggenschwiler et al., 2012) | TE/TR = 1.07/2400<br>TD<sub>1</sub>/TD<sub>2</sub> = 60/1800 | 4/11 | 2.0 | 256×256×144<br>(sagittal slab) | 490 | 2× GRAPPA<br>6/8×6/8 partial Fourier | 2.25 |
| fMRI 1.6 mm | 2D SMS GE-EPI<br>(Setsompop et al., 2012) | TE/TR = 23/1050 | 54 | 1.6 | 208×208×134<br>(84 axial slices) | 1924<sup>c</sup> | 2× GRAPPA, 4× SMS<br>7/8 partial Fourier | PD<sup>b</sup> |
| fMRI 0.8 mm | 2D SMS GE-EPI<br>(Setsompop et al., 2012) | TE/TR = 29/3520 | 81 | 0.8 | 192×192×116<br>(132 ax. slices) | 1226<sup>c</sup> | 3× GRAPPA, 3× SMS<br>6/8 partial Fourier | PD<sup>b</sup> |
| Ref. EPI distortion<br>correction 1.6 mm<sup>d</sup> | 2D SE-EPI | TE/TR = 40/7310 | 86 | 1.6 | 208×208×134<br>(84 axial slices) | 1924 | 2× GRAPPA<br>7/8 partial Fourier | 1.5 |
| Ref. EPI distortion<br>correction 0.8 mm<sup>d</sup> | 2D SE-EPI | TE/TR = 49/12170 | 90 | 0.8 | 192×192×116<br>(132 ax. slices) | 1226 | 3× GRAPPA<br>6/8 partial Fourier | 3.2 |

a. All protocols were defined with isotropic resolution; the value shown for each entry corresponds to the voxel width in all three directions.
b. PE: phase encoding; PD: paradigm-dependent (described in section 2.1.3).
c. Acquired with posterior-to-anterior PE direction.
d. SE-EPI with in-plane encoding parameters and under-sampling matched to the main fMRI protocol; comprised 3 volumes acquired in posterior-to-anterior PE direction, and 3 in the opposite sense.

**Supp. Table II.**
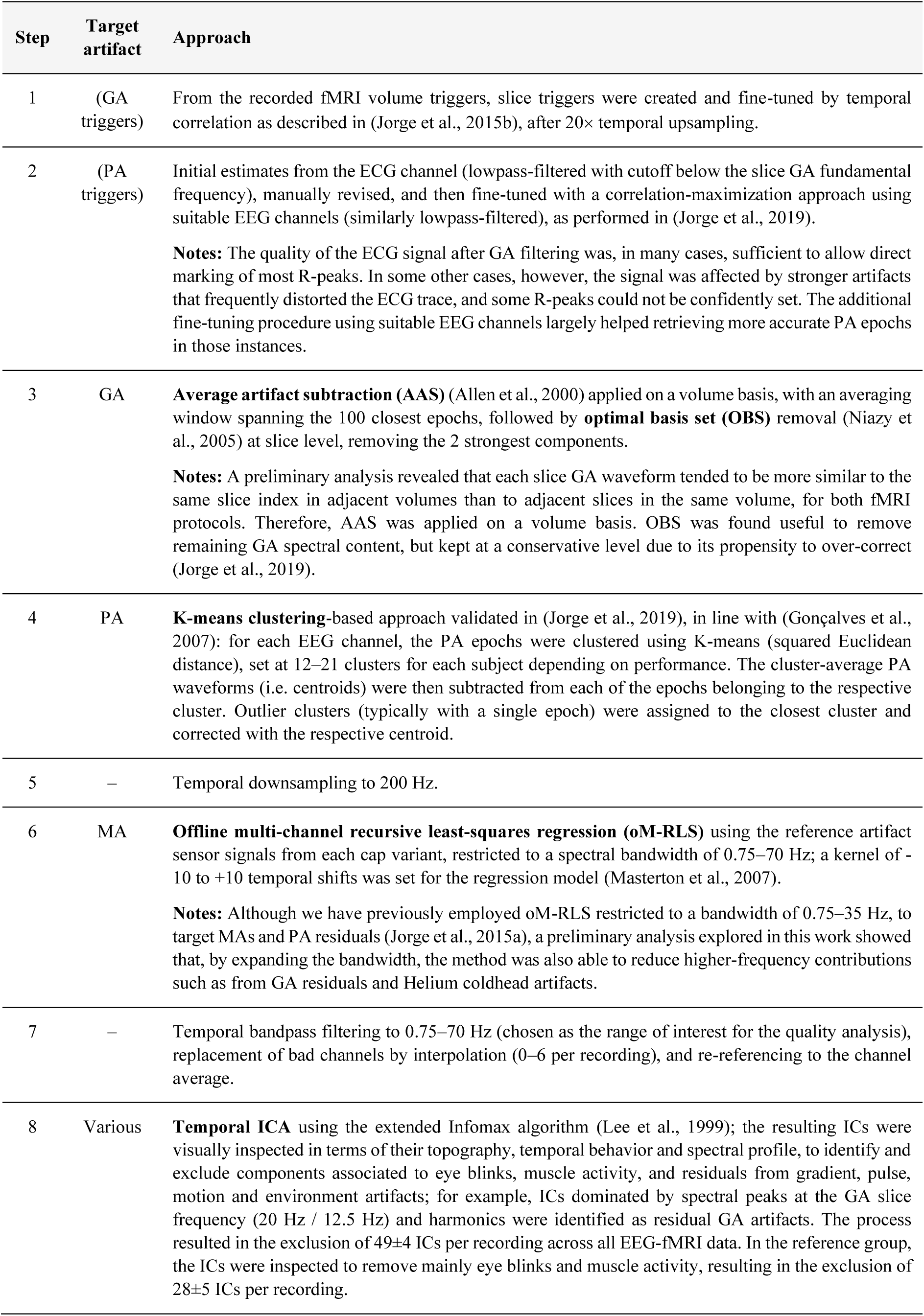
EEG artifact correction approach implemented for this work.

| Step | Target artifact | Approach |
| --- | --- | --- |
| 1 | (GA triggers) | From the recorded fMRI volume triggers, slice triggers were created and fine-tuned by temporal correlation as described in (Jorge et al., 2015b), after 20× temporal upsampling. |
| 2 | (PA triggers) | Initial estimates from the ECG channel (lowpass-filtered with cutoff below the slice GA fundamental frequency), manually revised, and then fine-tuned with a correlation-maximization approach using suitable EEG channels (similarly lowpass-filtered), as performed in (Jorge et al., 2019).<br>**Notes:** The quality of the ECG signal after GA filtering was, in many cases, sufficient to allow direct marking of most R-peaks. In some other cases, however, the signal was affected by stronger artifacts that frequently distorted the ECG trace, and some R-peaks could not be confidently set. The additional fine-tuning procedure using suitable EEG channels largely helped retrieving more accurate PA epochs in those instances. |
| 3 | GA | **Average artifact subtraction (AAS)** (Allen et al., 2000) applied on a volume basis, with an averaging window spanning the 100 closest epochs, followed by **optimal basis set (OBS)** removal (Niazy et al., 2005) at slice level, removing the 2 strongest components.<br>**Notes:** A preliminary analysis revealed that each slice GA waveform tended to be more similar to the same slice index in adjacent volumes than to adjacent slices in the same volume, for both fMRI protocols. Therefore, AAS was applied on a volume basis. OBS was found useful to remove remaining GA spectral content, but kept at a conservative level due to its propensity to over-correct (Jorge et al., 2019). |
| 4 | PA | **K-means clustering**-based approach validated in (Jorge et al., 2019), in line with (Gonçalves et al., 2007): for each EEG channel, the PA epochs were clustered using K-means (squared Euclidean distance), set at 12–21 clusters for each subject depending on performance. The cluster-average PA waveforms (i.e. centroids) were then subtracted from each of the epochs belonging to the respective cluster. Outlier clusters (typically with a single epoch) were assigned to the closest cluster and corrected with the respective centroid. |
| 5 | – | Temporal downsampling to 200 Hz. |
| 6 | MA | **Offline multi-channel recursive least-squares regression (oM-RLS)** using the reference artifact sensor signals from each cap variant, restricted to a spectral bandwidth of 0.75–70 Hz; a kernel of -10 to +10 temporal shifts was set for the regression model (Masterton et al., 2007).<br>**Notes:** Although we have previously employed oM-RLS restricted to a bandwidth of 0.75–35 Hz, to target MAs and PA residuals (Jorge et al., 2015a), a preliminary analysis explored in this work showed that, by expanding the bandwidth, the method was also able to reduce higher-frequency contributions such as from GA residuals and Helium coldhead artifacts. |
| 7 | – | Temporal bandpass filtering to 0.75–70 Hz (chosen as the range of interest for the quality analysis), replacement of bad channels by interpolation (0–6 per recording), and re-referencing to the channel average. |
| 8 | Various | **Temporal ICA** using the extended Infomax algorithm (Lee et al., 1999); the resulting ICs were visually inspected in terms of their topography, temporal behavior and spectral profile, to identify and exclude components associated to eye blinks, muscle activity, and residuals from gradient, pulse, motion and environment artifacts; for example, ICs dominated by spectral peaks at the GA slice frequency (20 Hz / 12.5 Hz) and harmonics were identified as residual GA artifacts. The process resulted in the exclusion of 49±4 ICs per recording across all EEG-fMRI data. In the reference group, the ICs were inspected to remove mainly eye blinks and muscle activity, resulting in the exclusion of 28±5 ICs per recording. |

